# Redefining CD4 T cell residency: Helper T cells orchestrate protective humoral immunity in the lung

**DOI:** 10.1101/2020.02.28.963280

**Authors:** Nivedya Swarnalekha, David Schreiner, Ludivine C Litzler, Saadia Iftikhar, Daniel Kirchmeier, Marco Künzli, Carolyn G King

## Abstract

Influenza is a severe and acute respiratory pathogen, and a significant cause for morbidity, particularly in young children and the elderly. Following influenza infection, clonally expanded T cells take up permanent residence in the lung where they are poised to rapidly respond to challenge infection. The non-circulating status of these tissue resident memory (TRM) cells makes them an attractive target for vaccination. While many studies have characterized CD8 TRM cells, less is known about the heterogeneity and protective capacity of CD4 TRM cells. Here we characterized the dynamics and transcriptional regulation of lung resident CD4 T cells to define a non-lymphoid signature that removes the bias created by the prevalence of Th1 helper cells during viral infection. We identified a novel population of long-lived T resident helper (TRH) cells that requires intrinsic Bcl6 expression for their differentiation. Although TRH cells also depend on B cells, they are generated independently of T follicular helper effector cells in the lymph node. In contrast to lung resident Th1 cells, TRH cells are tightly co-localized with B cells in inducible Bronchus Associated Lymphoid Tissue (iBALT). Deletion of Bcl6 in CD4 T cells prior to heterotypic challenge infection results in redistribution of CD4 T cells outside of iBALT areas and impaired local antibody production. These data highlight lung iBALT as a niche for the homeostasis and survival of TRH cells, and further suggest that vaccination strategies to selectively induce TRH cells can improve protective immunity in the tissue.

## Main

Seasonal influenza epidemics are a major cause of global morbidity and mortality. Although annually administered influenza vaccines are among the most widely used in the world, vaccine-elicited neutralizing antibodies offer poor protection against new influenza strains^1^. In contrast, there is evidence that prior influenza infection can accelerate viral clearance after heterotypic infection in both mice and humans^2–6^. Emerging data suggests that the targeted generation of CD4 memory T cells recognizing conserved epitopes from internal viral proteins may form the basis of a universal influenza virus vaccine^7, 8^. CD4 memory T cells are induced following immunization or infection and can be recalled to generate secondary effectors during a challenge infection. Several subsets of CD4 memory cells have been described, including central memory (TCM) and effector memory (TEM) cells which circulate through secondary lymphoid and non-lymphoid tissues^9^. More recently, tissue resident memory (TRM) cells that persist in barrier tissues such as lung and skin have been described^10^. Although CD4 T cells actually outnumber CD8 T cells in barrier tissues, the majority of studies have focused on the requirements for CD8 TRM cell differentiation. In addition, although CD4 T cells are renowned for their substantial plasticity during immune responses, less is known about diversification within the CD4 TRM cell compartment^11–16^.

Influenza infection induces the differentiation of CD4 TRM cells in the lung, where they are maintained in an antigen and inflammation independent manner^17^. Following a lethal re-challenge, influenza specific CD4 TRM cells rapidly produce effector cytokines and promote both viral clearance and host survival^7^. Lung CD4 TRM cells can also be induced by mucosal vaccination and were shown to mediate superior protection following heterologous infection, highlighting their potential as a universal vaccine target^18^. Influenza-specific CD4 TRM cells are generally characterized as Th1-like, with the capacity to produce both IFNγ and IL-2^7, 19^. However, IL-2 deficient memory CD4 T cells were recently shown to provide superior protection compared to wild-type memory cells, an outcome that correlated with decreased inflammation and host pathology during re-challenge^20^. These data suggest that protection mediated by CD4 TRM cells is not strictly dependent on their ability to produce effector cytokines, and that a balanced secondary response is likely to involve the recruitment and coordination of distinct and specialized CD4 TRM cell subsets^21^.

Heterogeneity within the non-antigen specific CD4 TRM cell compartment was recently addressed in a study that reported enrichment of genes associated with the TNFRSF and NFKB pathways in barrier T cells compared to T cells isolated from their respective draining lymphoid compartments^11^. The residency signature derived from this dataset, however, does not take into consideration tissue versus lymphoid organ differences between distinct T cell subsets. Accordingly, this approach tends to over-represent genes associated with type 1 helper T cells such as *Klrg1*, *Itgae*, *Id2*, and *CXCR6*, which may comprise a poorer definition of residency for other Th cell subsets.

Consistent with this idea, the authors reported the presence of a stronger TRM phenotype in lymphoid Th1 memory cells compared to lymphoid TFH cells which they attributed to the relative ability of these cells to adapt to barrier tissues. On the other hand, it is well appreciated that TFH cells share many surface markers and molecular dependencies with TRM cells, including high expression of PD1, P2X7R, CD69 and ICOS, and a requirement for S1PR1 and KLF2 downregulation to develop^10, 22–25^. In support of this idea, TFH memory cells isolated from the spleen after LCMV infection were recently demonstrated to have a partially overlapping transcriptional signature with TRM cells, consistent with the non-circulating nature of both subsets^26^. In contrast, another study addressing the relationship between TRM cells present in secondary lymphoid organs and TFH memory cells reported more distinct transcriptional profiles of these subsets^12^. However, in this case the authors focused on gene expression in T cell receptor (TCR) transgenic cells that have relatively impaired TFH memory cell generation compared to polyclonal antigen specific cells^26^. In addition, although the authors reported a strong overlap between CD4 and CD8 TRM signatures after LCMV infection, these signatures are likely to be skewed by the induction of genes expressed by cytotoxic and type 1 cytokine producing cells. Thus, an open question is whether different tissue resident Th cell subsets express conserved or distinct residency signatures compared to their lymphoid counterparts.

In this study we characterized the dynamics and diversification of polyclonal CD4 T cells present in the lung and draining LN after influenza, using the resolution provided by scRNAseq to decouple residency and functional signatures between these two compartments. Our analyses of influenza-specific CD4 T cells reveal striking heterogeneity within the lung compartment comprised of two broad subsets which we designate tissue resident helper (TRH) and tissue resident memory 1 (TRM1) cells.

TRH cells persist stably and depend on expression of the transcription factor Bcl6 for their differentiation. TRH are tightly localized together with B cells in the lung and require ongoing antigen presentation for their maintenance. Importantly, CD4 T cell intrinsic deletion of Bcl6 at late time points after primary infection impairs the local humoral response upon reinfection. These data identify a novel TRH subset that may be a rational target to drive potent and protective immunity in the lung mucosa^11, 12^.

## Results

### Residency signature in CD4 T cells is biased toward Th1 memory cells

To examine influenza-specific CD4 memory T cell populations, we performed scRNA-seq on memory CD4 T cells from the lung and lung-draining mediastinal LN. Virus specific T cells were detected by tetramer staining for IA^b^:NP^311–325^ and were largely protected from intravascular antibody staining for CD45 (Fig. 1a, Supplementary Fig. 1a). Treatment of mice with fingolimod (FTY720) to block the egress of memory cells from lymphoid organs did not alter the total number of NP+ CD4 T cells isolated from the lung, indicating that this population can be sustained without input from the circulation at this time point (Supplementary Fig. 1b, c). Following quality control, normalization and dimension reduction, the transcriptomes of lung and LN T cells were clearly distinct, and an expected small population of putative circulating cells identified by elevated expression of *S1pr1*, *Il7r* and *Bcl2* was also detected in the lung (Fig. 1b, c, Supplementary Table 1a). Differential expression analysis between lung and LN cells confirmed enrichment of previously reported CD4 residency genes (e.g. *Crem*, *Vps37b*, *Rora*, *Ramp3*, *Tnfrsf18*) in the lung (Fig. 1c)^11, 12^. To determine if the residency signature was conserved in another viral infection model we analyzed GP66 specific memory T cells from the liver and spleen of LCMV infected mice (Fig 1d). Lung specific genes from influenza were broadly enriched in liver cells from LCMV and vice versa (Supplementary Fig. 1d). To better define a conserved residency signature, we took the intersection of genes differentially enriched in non-lymphoid tissues (NLT) from both viral infections (Fig. 1e). This “pseudo-bulk”, tissue-level analysis demonstrates an apparent bias in NLT toward a Th1 phenotype, with *Ccl5*, *Crip1*, *Vim*, *Nkg7*, *Id2* and *Cxcr6* among the top shared genes (Fig. 1e, f). Unsurprisingly, a multi-tissue signature for CD8 TRM was also enriched in lung and liver, consistent with similarities between the transcriptional regulation of cell identity among type I cytokine producing CD4 and CD8 cells (Fig. 1g). Conversely, TFH signatures including genes such as *Tcf7*, *Izumo1r*, *Dennd2d*, *Shisa5*, *Rgs10* and *Cxcr5* were enriched in lymphoid tissues (LT), in line with the conception of TFH as an SLO-resident population (Fig. 1e, f)^22^. The presence of Th subset-specific genes in both NLT and LT cells reported here and elsewhere calls for a refinement of the residency signature that separates function from niche^11, 12^.

**Fig. 1.**
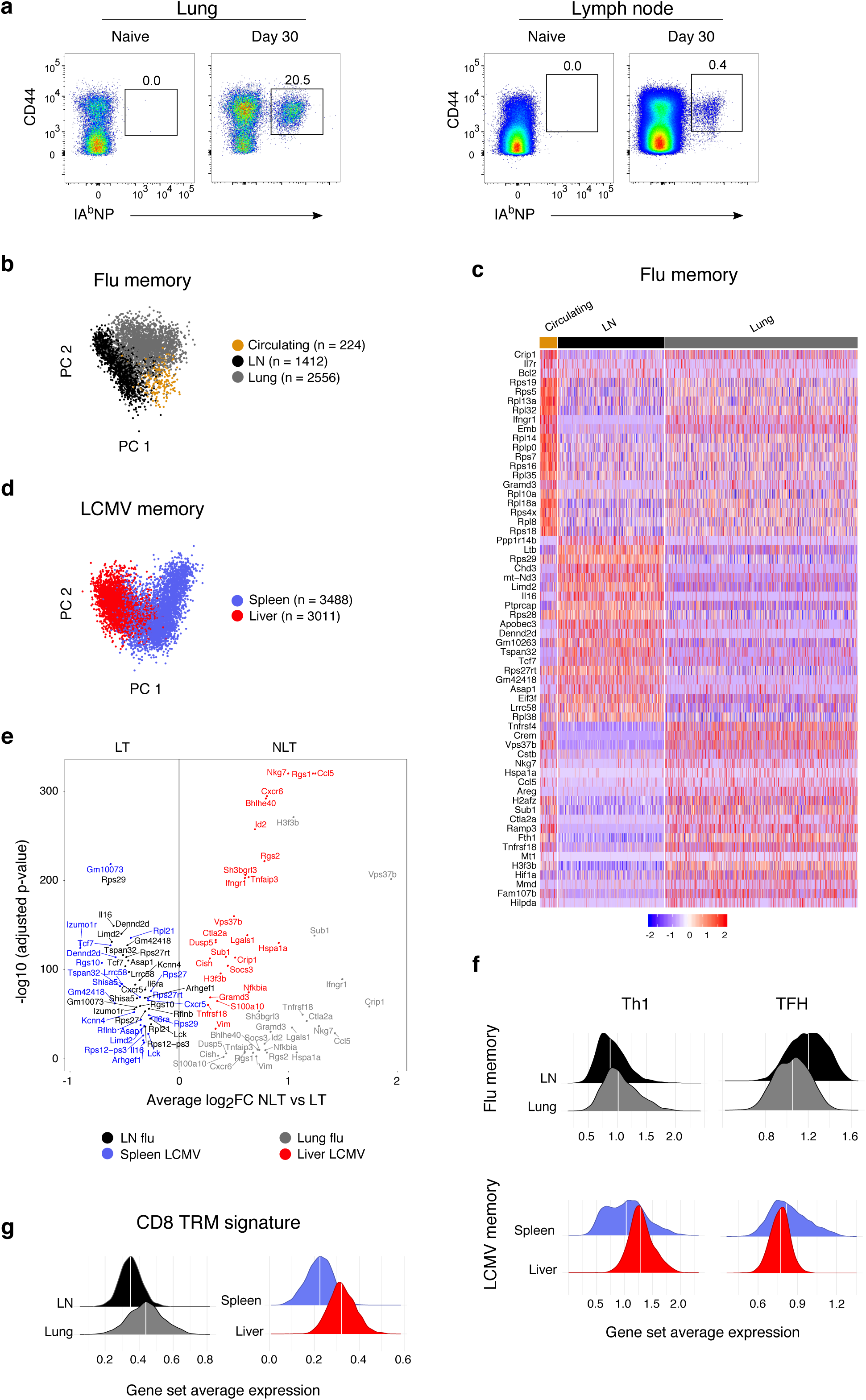
Residency signature in CD4 T cells is biased toward Th1 memory cells. **a**, NP-specific CD4 T cells in lung and mLN of naïve and PR8 memory mice. **b,** PCA showing LN and lung samples with putative circulating cluster from lung^26^. **c,** Heatmap showing scaled, centered single cell expression of top 20 genes sorted according to cluster average log_2_FC, adjusted P value < 0.05. **d,** PCA showing spleen and liver samples, LCMV infection. **e,** Differentially expressed genes discriminating lung from LN (flu) or liver from spleen (LCMV). P values of 0 assigned the lowest non-zero P value. **f,** Log-normalized average expression of TFH and Th1 memory signatures^26^. **g,** Log-normalized average expression of combined CD8 residency signature from multiple tissues^61, 62^.

### Heterogeneity in the lung: pan residency versus Th-subset specific residency

We next investigated heterogeneity specifically within the NP-specific lung CD4 T cell compartment. Unsupervised hierarchical clustering classified the cells into four populations (Fig. 2a). Notably, we found clear evidence of a TFH-like phenotype in the lung: cluster 3 showed increased expression of *Sostdc1*, *Sh2d1a*, *Ppp1r14b*, *Rgs10, Id3* and *Tcf7* (Fig 2b, Supplementary Table 2a). Although this TFH-like cluster was lower in many of the genes characterizing residency compared to the other lung clusters, expression was still clearly higher compared to cells in the LN (Supplementary Fig. 2a). Cells in cluster 2 had a Th1 phenotype including enrichment for *Selplg*, *Nkg7*, *Ccl5, Id2* and *Cxcr6*. The similarity of these clusters to known T-helper subsets was further confirmed by scoring each cell according to published TFH and Th1 memory gene sets (Fig. 2c). Elevated transcription of *S1pr1*, *Il7r* and *Bcl2* suggested that cluster 4, distinct in principal component 2 and further distinguished by high ribosomal protein content, contained circulating cells recovered during lung processing. Strikingly, while cluster 1 was most similar to the TFH-like cluster, it had a distinct profile characterized by enrichment for *Hif1a*, *Areg*, *Tnfrsf4* and *Tnfsf8* and also expressed more *Nr4a3,* suggesting ongoing MHC-II engagement and activation of the calcineurin/NFAT pathway (Fig. 2a-c, Supplementary Fig. 2b)^27^. Cells in cluster 1 were also enriched for *Rora*, which is reportedly induced by extrinsic signals in the microenvironment and plays a role in negatively regulating lung inflammation during infection^28^.

**Fig. 2.**
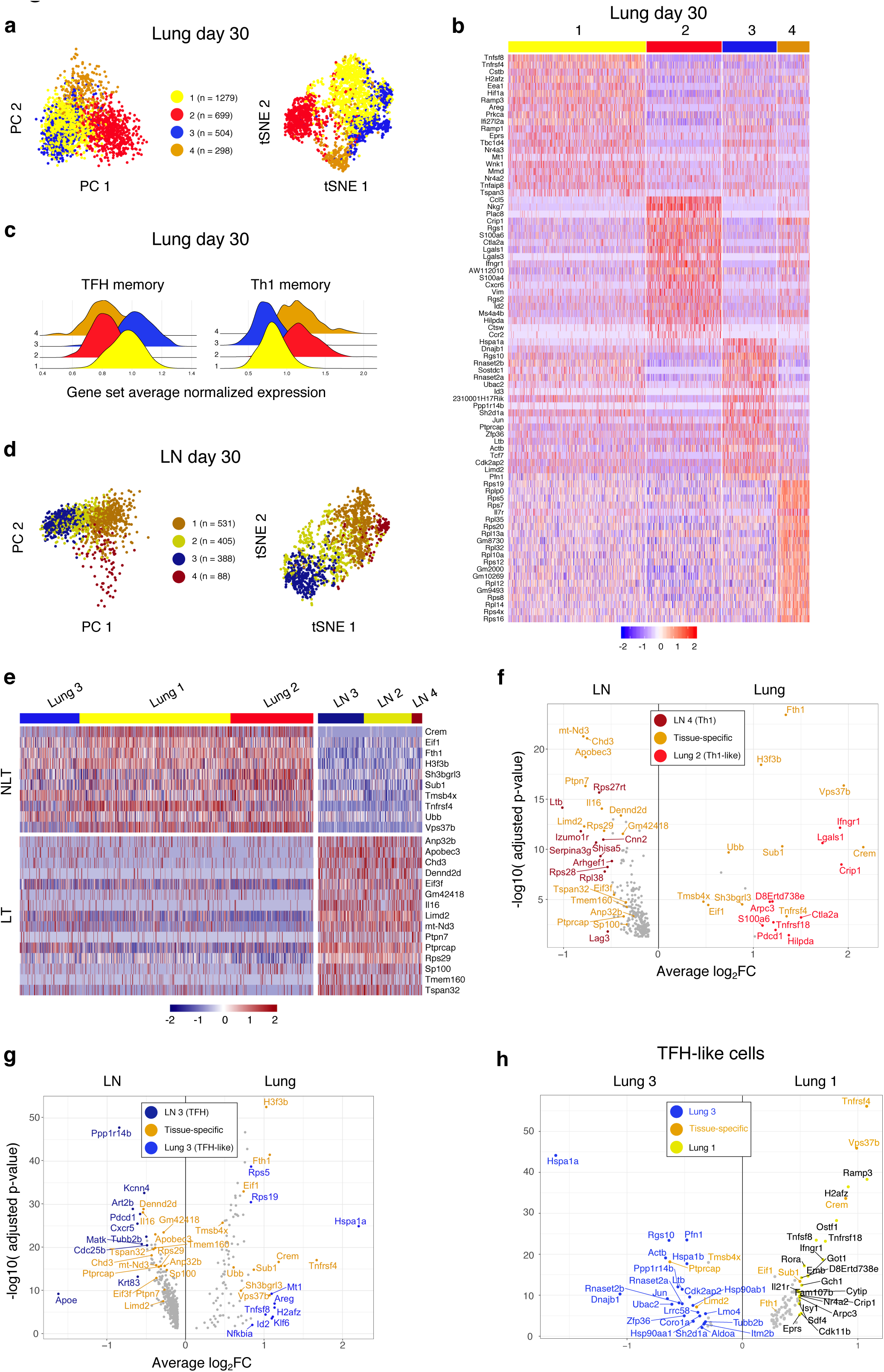
Heterogeneity in the lung: pan residency versus Th-subset specific residency. **a,** Unsupervised hierarchical clustering using Ward’s method of lung cells only: PCA and tSNE dimension reductions. **b,** Heatmap showing scaled, centered single cell expression of top 20 genes sorted according to lung cluster average log_2_FC, adjusted P value < 0.05. **c,** Log-normalized average expression of TFH and Th1 memory signatures^26^. **d,** Unsupervised hierarchical clustering using Ward’s method of LN cells only: PCA and tSNE dimension reductions. **e,** Heatmap of scaled, centered single cell expression showing conserved genes for NLT and LT. Genes included are differentially expressed in the tissue in all three of the non-circulating cluster pairs. Adjusted P value < 0.05. **f,** Differentially expressed genes discriminating LN Th1 cells from lung Th1-like cells with reference to conserved tissue-specific signature. Adjusted P value < 0.05. **g,** Differentially expressed genes discriminating LN TFH cells from lung TFH-like cells with reference to conserved tissue-specific signature. Adjusted P value < 0.05. **h,** Differentially expressed genes discriminating lung TFH-like cluster 3 from lung TFH-like cluster 1 with reference to conserved tissue-specific signature. Adjusted P value < 0.05.

We next focused our analysis on NP-specific CD4 T cells from the draining LN, cutting the resulting hierarchical clustering tree to four clusters (Fig. 2d). These clusters were classified similarly to those found in the lung, with a notably small Th1 cluster 4 expressing *Nkg7*, *Ccl5*, *Id2* and *Cxcr6* and tracking with ribosomal protein-rich cluster 1 along PC1; strongly TFH cluster 3 with high expression of *Pdcd1*, *Sh2d1a*, *Sostdc1*; and TFH-like cluster 2 enriched for *Izumo1r*, *Tox*, and *Hif1a* (Supplementary Fig. 2c, Supplementary Table 2b). As with the lung data, we scored LN cells using TFH and Th1 gene sets to further verify this classification (Supplementary Fig. 2d). To arrive at a less biased signature for lung residency, we performed differential expression analysis on each of the most phenotypically similar cluster pairs between lung and LN, for example, comparing LN TFH with TFH-like lung cells and LN Th1 with Th1-like lung cells. The final conserved residency signature consisted only of genes that were differentially enriched in all of the non-circulating lung clusters over their LN counterparts (Fig. 2e). This enrichment was confirmed in our LCMV data set as well as other recently published TRM data (Supplementary Fig. 2e-g). Subsequent removal of these conserved residency genes allowed us to identify Th-subset specific genes differentially expressed between tissue and lymphoid compartments. Th1-like cells in the lung were enriched over their LN counterparts for genes including *Ctla2a*, *Crip1*, *Lgals1*, *Tnfrsf18* and *Infgr1* while TFH-like lung cells exhibited higher expression of *Mt1*, *Hspa1a*, *Klf6*, *Tnfsf8*, and *Areg* (Fig. 2f, g, Supplementary Table 2c, d). In addition, we compared the two TFH-like clusters in the lung and found a more lymphoid signature in cluster 3 and enrichment of the residency signature in cluster 1, suggesting functional diversification of TFH-like cells in the lung (Fig. 2h, Supplementary Table 2e).

To gain further insight into the transcriptional regulation of distinct resident CD4 T cell subsets, we next assessed transcription factor (TF) activity using single-cell regulatory network inference and clustering (SCENIC)^29^. This approach identified active regulatory activity in TFs such as Hif1a, Crem, Fosl2, and known Th1-related factors Prdm1, Runx2 and Runx3 (Supplementary Fig. 2h). Cluster-specific analysis revealed that the activity of Prdm1, Runx2 and Runx3 was limited to Th1-like clusters while Hif1a activity was focused primarily in lung cluster 1, mirroring gene expression of *Hif1a*. Bcl6 regulatory activity was detected in lung TFH-like cluster 3 and LN TFH cluster 3, along with Cebpa, which is reported to inhibit both IFNγ production and TCR-driven proliferation (Supplementary Fig. 2i)^30, 31^. Interestingly, Foxp3-associated regulatory activity was enriched in lung over LN although there was only minimal expression of *Foxp3* transcript in the data set and Foxp3+ cells were not detected by FACS in the antigen-specific compartment (Supplementary Fig. 2j, k). In summary, the scRNA-seq data gathered here constitute an extensive picture of CD4 T cell memory to influenza and disentangle residency from helper function in order to highlight not only conserved differences between NLT and LT, but also intra-tissue heterogeneity and functional diversification of lung resident T cells.

### Phenotypic characterization of resident CD4 T cell subsets

To determine if the heterogeneity detected by scRNAseq was also mirrored at the protein level, we examined the phenotype of NP-specific CD4 T cells using flow cytometry. At day 30 after infection, NP-specific CD4 T cells in the lung could be divided into two major subsets with reciprocal expression of FR4 and PSGL1 (Fig. 3a). Following viral clearance and T cell contraction, the number of cells falling within these two T resident cell subsets remained stable to at least 120 days after infection (Fig. 3b). Treatment with FTY720 at day 30 after infection did not change the number of cells in either subset, indicating that both populations are resident (Supplementary Fig. 3a).

**Fig. 3.**
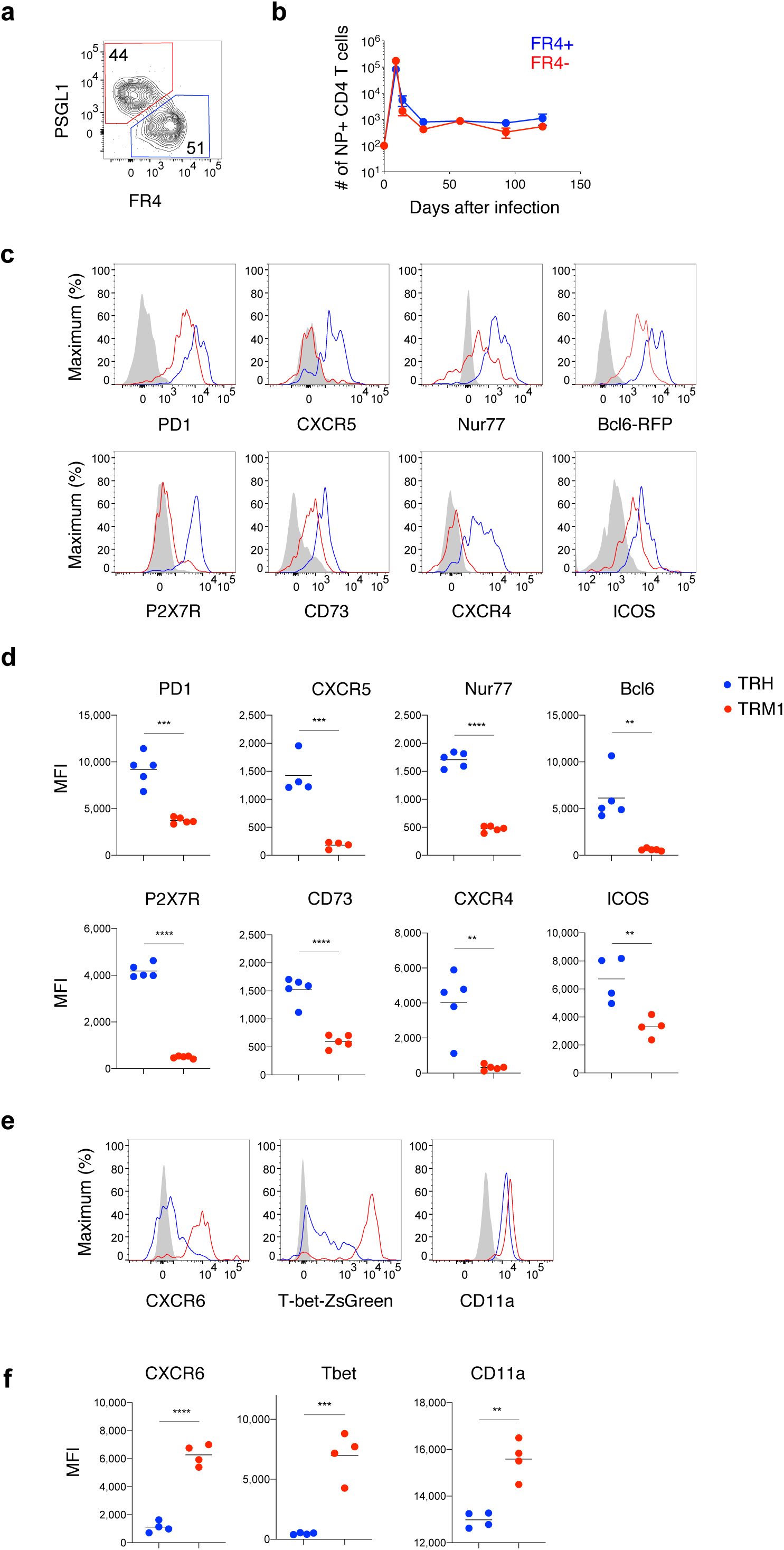
Phenotypic characterization of resident CD4 T cell subsets. **a,** Identification of TRM1 (red) and TRH (blue) subsets in NP-specific resident CD4 T cells. **b,** Total numbers of FR4^+^ and FR4^-^ NP-specific T cells over time. **c-f,** Histograms **c, e** and MFI **d, f** of indicated phenotypic markers in TRH (blue), TRM1 (red) and naïve CD4 T cells (gray). Data are represented as the mean ± s.e.m (n=5, representing 2 experiments) in **b.** Thin lines represent mean in **d** and **f** (n=4-5, representing 2 experiments). The significance was determined by unpaired Student’s t-test. P values: *P<0.05, **P<0.01, ***P<0.001, ****P<0.0001.

FR4^hi^PSGL1^low^ (hereafter T resident helper, TRH) cells expressed many markers associated with TFH effector cells including PD1, CXCR5, Nur77, Bcl6, P2X7R, CD73, CXCR4 and ICOS (Fig. 3c, d). FR4^lo^PSGL1^hi^ (hereafter TRM1) cells expressed lower levels of TFH-associated markers, but higher levels of markers associated with Th1 cells including CXCR6, T-bet and CD11a (Fig. 3e, f). Importantly, adoptively transferred OT-II T cell receptor transgenic cells activated by infection with PR8-OVA2 generated mostly TRM1 cells (Supplementary Fig. 3b, c). These data are consistent with earlier reports using TCR transgenic strains that primarily identified Th1-like memory CD4 T cells in the lung after influenza infection. Taken together, scRNAseq and flow cytometry confirm the presence of TRM1 (Th1-like) and TRH (TFH-like) cells in the lung and underscore the importance of studying polyclonal CD4 T cell responses to understand T-dependent immunity.

### Progressive differentiation of resident CD4 T cell subsets

To further investigate the kinetics of TRM1 and TRH cell differentiation in the lung, we next characterized the phenotype of NP-specific CD4 T cells at multiple time points after infection. At day 9, the majority of CD4 T cells in the lung had high expression of PSGL1, bimodal expression of FR4 and were negative for TFH markers CXCR5 and PD1 (Fig. 4a, b). By day 14 after infection, a small proportion of TRH cells emerged in the lung, followed by a clear separation of TRM1 and TRH cell subsets by day 30 (Fig. 4a). In contrast to the lung, fully differentiated NP-specific TFH effector cells (CXCR5+PD1+) were clearly detected in the mediastinal LN at day 9 (Fig. 4b). To understand if LN TFH effectors give rise to TRH cells, we began continuously treating mice with FTY720 between days 9 and 30 to prevent the migration of LN-primed TFH cells into the lung. At day 30 after infection, the number of TRH cells in FTY720 treated mice was equivalent to the number of TRH cells in PBS treated mice, indicating that full differentiation of lung TRH cells can be independently completed in the lung (Fig. 4c).

**Fig. 4.**
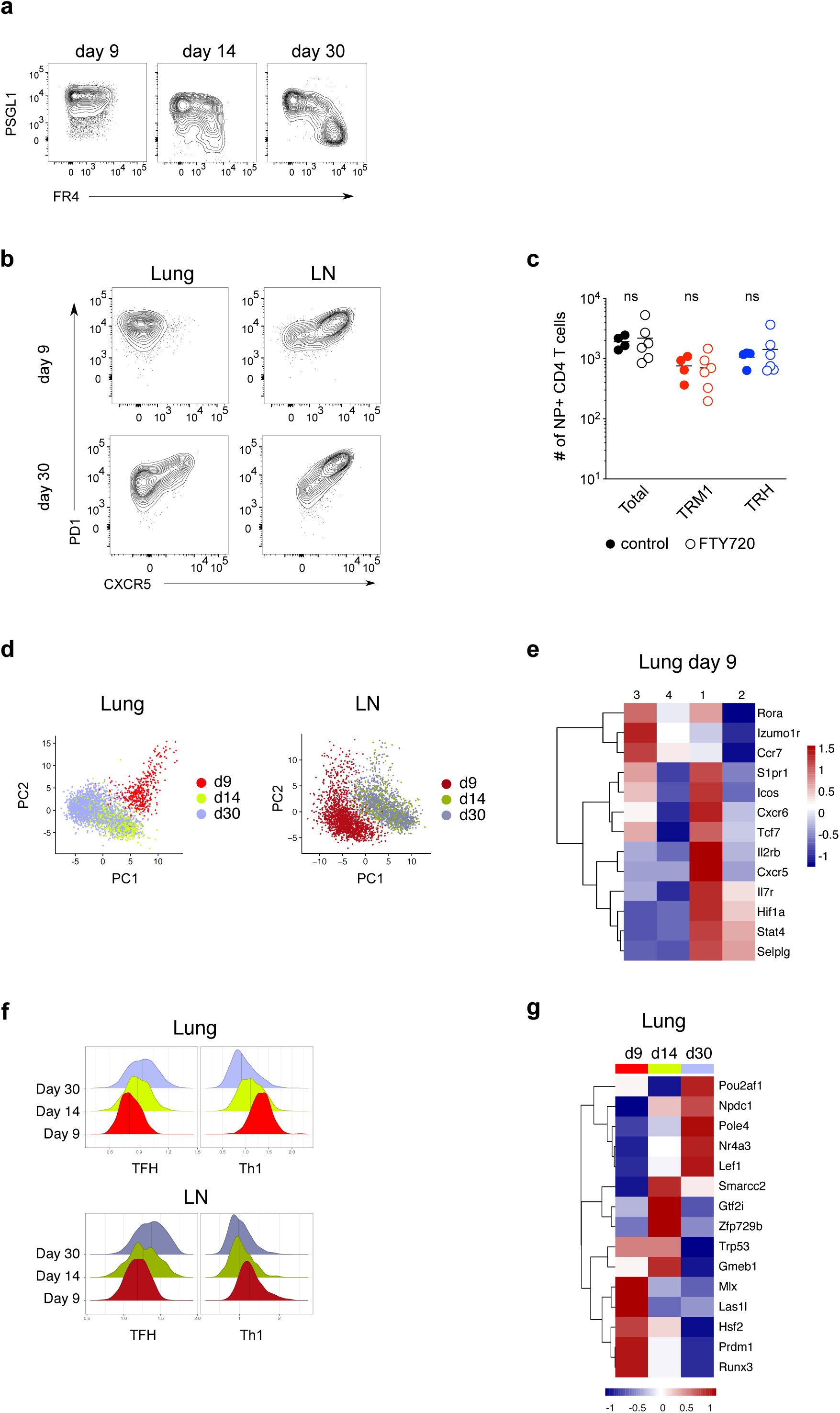
Progressive differentiation of resident CD4 T cell subsets. **a,** FR4 and PSGL1 expression on NP-specific tissue resident CD4 T cells at indicated time points. **b,** CXCR5 and PD1 expression on NP-specific CD4 T cells in lung and mLN at indicated time points. **c,** NP-specific total CD4 T cells, TRM1 and TRH cell numbers in control and FTY720 treated mice. Data in **a, b, c** represent 2 experiments with n=4-6 mice. Statistical significance was determined by unpaired Student’s t-test. Thin lines represent mean. p<0.05 is considered significant. **d,** PCA showing time point by tissue. **e,** Scaled, centered average expression of select genes in lung at day 9. **f,** Log-normalized average expression of published TFH and Th1 signatures by time point and tissue^26^. **g,** Scaled, centered average area under curve (AUC) calculated with SCENIC showing top 5 differentially active transcription factors in lung by time point. Adjusted P-value < 0.01.

To investigate the transcriptional regulation of resident T cell generation, we expanded the scRNA-seq analysis to include NP-specific CD4 T cells from days 9 and 14 after infection. Cells were clearly discriminated by time point and tissue in reduced dimensions, with day 9 cells appearing more distinct compared to days 14 and 30 (Fig. 3d, Supplementary Fig. 4a). Day 9 lung cells exhibited the lowest per cell gene counts of any sample, which may reflect their susceptibility to apoptosis under inflammatory conditions. Nevertheless, hierarchical clustering revealed a group of cells (cluster 1) enriched for *Stat4, Il7r, Icos, S1pr1,* and *Selplg*, which may represent recent immigrants from the lymph node (Fig. 4e). Consistent with the FACS analysis, we observed an enrichment of Th1-like characteristics at day 9 while TRH cells were more prominent at day 30 (Fig. 4f). To investigate the expression of *Izumo1r* and *Selpg*, encoding the proteins FR4 and PSGL1, respectively, we used imputed values, replacing zeros with values estimated from similar cells. In parallel with FACS expression of these markers, day 9 lung cells were *Selplg*^hi^ and *Izumo1r*^lo^, while *Selplg*^lo^ and *Izumo1r*^hi^ T cells emerged at day 14 and were increased by day 30 (Supplementary Fig. 4b). In contrast, the majority of cells in the LN were Izumo1r^hi^ at all time points. Transcription factor activity according to SCENIC analysis tracked the trend from Th1-like at day 9 (Runx2, Runx3, Prdm1) to TFH-like at day 30 (Lef1, Pou2af1) (Fig. 4g)^32, 33^. In addition, a consistent subset of genes associated with lung residency (*Tnfrsf4*, *Vps37b*, *Fth1*, *Crem*) was detected by differential expression analysis between lung and LN at each separate time point (Supplementary Fig. 4c). Other genes associated with residency at day 30 (*Gch1, Hif1a, Selenok, Prkca)* appeared more gradually, potentially reflecting the dynamics of signal access in the tissue microenvironment and the relatively late development of TRH compared to TRM1 cells (Supplementary Fig. 4d). In summary, high expression of PSGL1 by the majority of CD4 T cells at day 9 likely reflects their recent migration into the inflammatory lung environment; this time point is dominated by TRM1 cells while TRH cells emerge gradually at later time points.

### TRH cell generation requires B cells and T cell intrinsic Bcl6

Given the phenotypic and transcriptional similarities between lung TRH cells and lymphoid homing TFH cells, we hypothesized that TRH cells may depend on B cells and intrinsic Bcl6 expression for their differentiation^23, 34^. To test a requirement for B cells, we injected mice with anti-CD20 one week prior to influenza infection. Thirty days after influenza infection, both the proportion and number of TRH cells were decreased in the lungs of B cell depleted mice, correlating with a strong reduction in resident and circulating B cell numbers in the lung (Fig. 5a-c, Supplementary Fig. 5a). To assess the contribution of T cell intrinsic Bcl6 expression we generated radiation bone marrow chimeras with a mixture of wild type and Bcl6^flox/flox^ x CD4^Cre^ (Bcl6Δ^CD4^) bone marrow. Thirty days after influenza infection, Bcl6 deficient CD4 T cells were incapable of differentiating into lung TRH cells, demonstrating their dependence on intrinsic Bcl6 expression (Fig. 5d). In the absence of Bcl6, the proportions of TRM1 cells in the lung and non-TFH cells in the draining LN were also significantly reduced compared to wildtype cells, suggesting that Bcl6 also plays a role in the expansion of these cells (Fig. 4e, Supplementary Fig. 5b, c)^35^. Taken together, these data indicate that similar to lymphoid TFH cells, TRH cells in the lung require B cells and T cell intrinsic Bcl6 expression for their generation.

**Fig. 5.**
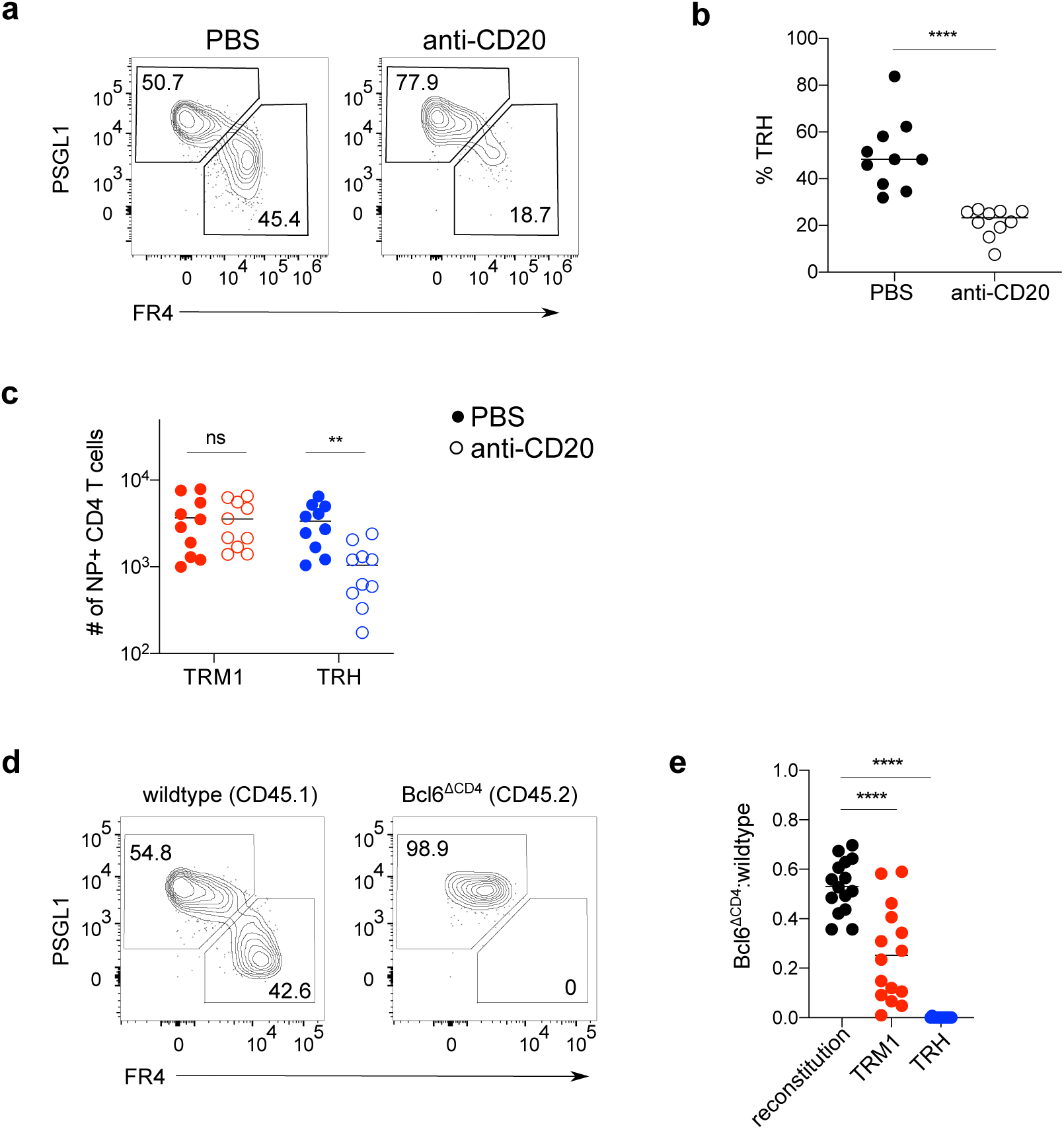
TRH cell generation requires B cells and T cell intrinsic Bcl6. **a, b** NP-specific T cells (**a)** and frequency of TRH (**b)** from control and B cell depleted mice. **c,** Total number of NP-specific TRM1 and TRH from control and B cell depleted mice. **d,** NP-specific T cells in control and Bcl6-deficient subsets. **e,** Bcl6Δ^CD4^:control ratio in CD4 T cells after reconstitution from blood (pre-infection), TRM1 and TRH. Thin lines represent mean in **b**, **c** (n=10, pooled from 2 experiments) and **e** (n=15, pooled from 3 experiments). Significance was determined by unpaired Student’s t-test. P values are as follows: *P<0.05, **P<0.01, ***P<0.001, ****P<0.0001.

### CD4 TRH cells localize in iBALT

Influenza infection leads to the development of ectopic clusters of B cells and T cells in the lung, known as inducible bronchus-associated lymphoid tissue (iBALT), that can persist to at least 100 days after infection^36, 37^. The formation of iBALT supports primary T and B cell proliferation and local antibody production and is thought to provide a specialized niche for the reactivation of lung immunity during challenge infection. Given their dependence on B cells, we hypothesized that TRH cells may localize together with B cells in iBALT. To examine this, we assessed the spatial distribution of TRM1 and TRH cells in the lungs of influenza infected T-bet and Bcl6 reporter mice, respectively. Using flow cytometry, we first confirmed that infection-induced polyclonal CD4 T cells in the lung largely phenocopy NP-specific TRM1 and TRH cells (Supplementary Fig. 6a). Gating on the minimal markers CD4 and T-bet, or CD4 and Bcl6, revealed that over 85% of these cells fall within the TRM1 or TRH compartments, respectively (Supplementary Fig. 6b, c). We next stained lung sections from T-bet or Bcl6 reporter mice with CD4 and B220 to identify iBALT regions by spinning disk confocal microscopy (Fig. 6a). T cell localization was quantified using two separate pipelines: one manually guided and one leveraging machine learning (Fig. 6b-d). The number of Tbet^hi^ or Bcl6^hi^ CD4 cell objects inside and outside of the B cell clusters was calculated after normalization by iBALT and tissue slice volumes. Bcl6 ^hi^ CD4 cells were enriched inside of iBALT areas compared to Tbet ^hi^ CD4 cells, while Tbet ^hi^ CD4 cells were more prevalent outside of iBALT (Fig 6c, d). The preferential co-localization of TRH cells and B cells in the lung indicates that iBALT may provide a homeostatic niche for TRH cell survival.

**Fig. 6.**
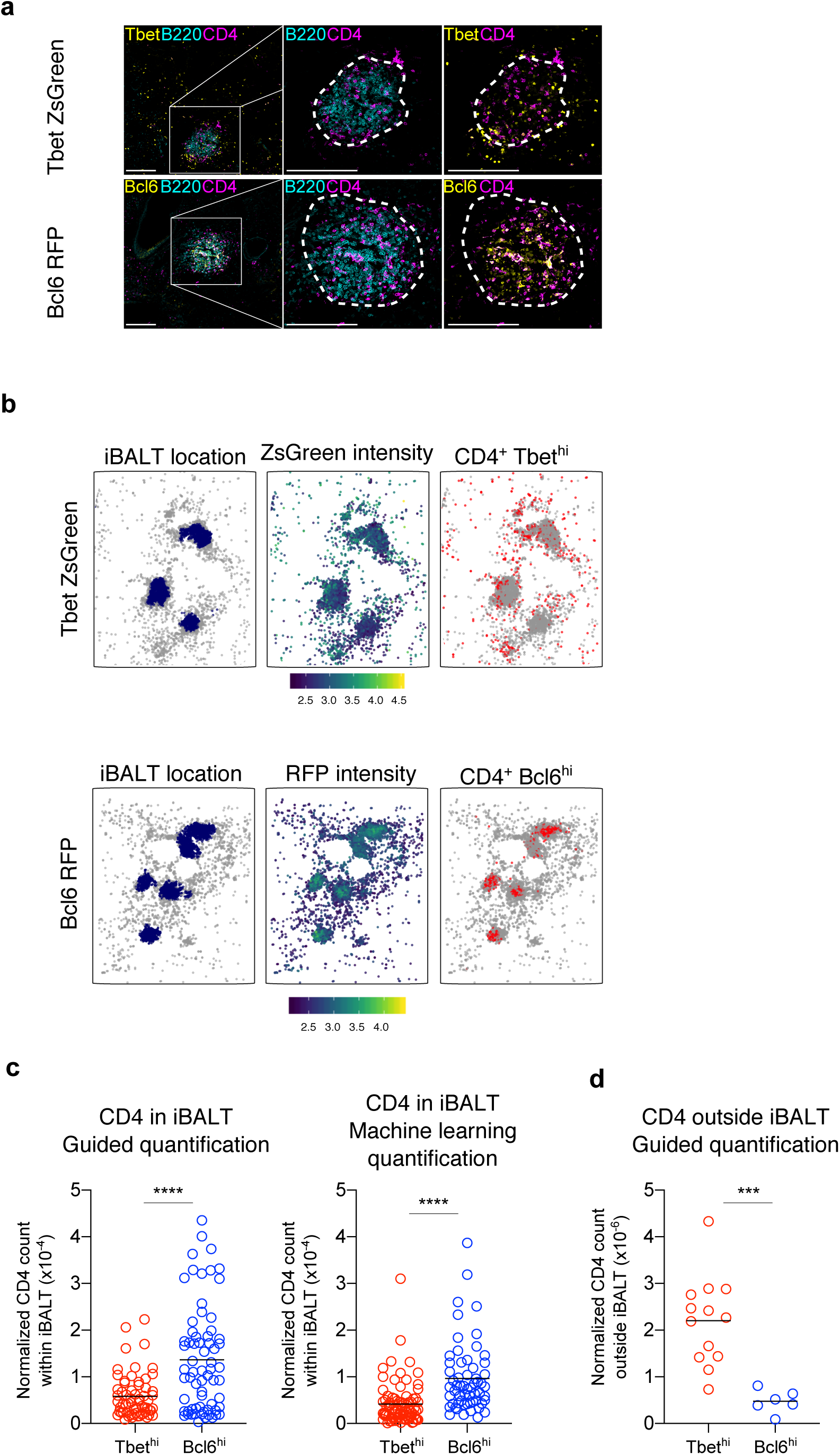
CD4 TRH cells localize in iBALT. **a,** Representative X40 immunofluorescence confocal images from Tbet ZsGreen and Bcl6 RFP mice, 30-60 days post-infection, stained with the indicated markers. Scale bars: 200um. **b,** Quantified representations of segmented CD4+ objects, showing iBALT location, log mean transcription factor intensity per object, and location of Tbet^hi^ (above) or Bcl6^hi^ (below) objects. **c,** Count of Tbet^hi^ and Bcl6^hi^ CD4 T cell objects inside B220+ iBALT clusters, normalized by iBALT volume, analyzed by Mann-Whitney-Wilcoxon test. Segmentation performed in Imaris (left) or with machine learning algorithm (right). Each dot represents one iBALT. **d,** Count of Tbet^hi^ and Bcl6^hi^ CD4 T cell objects outside B220+ iBALT clusters, normalized by tissue volume less iBALT volume, analyzed by Mann-Whitney-Wilcoxon test. Segmentation performed in Imaris. Each dot represents one tissue slice.

### Maintenance of TRH cells requires antigen presentation

Influenza infection has been shown to induce a depot of viral antigen in the lung which can be transported to and presented in the draining LN to support CD4 memory T cell accumulation^38^. High expression of Nur77 by TRH cells (Fig. 3c, d) suggested that they might be responding to ongoing antigen presentation in the lung.

To determine if TRH cells are actively proliferating at late time points, we administered bromo-2-deoxyuridine (BrdU) in the drinking water of influenza infected mice at late time points after influenza infection. Given the low number of NP-specific T cells that can be recovered from the lung after intranuclear staining for BrdU, we analyzed both NP+ as well as CD4+44+ T cells that were negative for intravascular staining with CD45. Following BrdU treatment, CD4 TRM cells took up significantly less BrdU compared to “circulating” lung CD4 T cells that stained positively for i.v. anti-CD45 (Fig. 7a, b). Comparison of BrdU uptake by TRH and TRM1 cells, however, revealed similar levels of proliferation between these two subsets, despite higher expression of Nur77 by lung TRH cells (Fig. 7c). To further determine whether antigen presentation is required to sustain resident CD4 T cells, we generated MHC-II^flox/flox^ x UBC^Cre-ERT2^ mice (MHC-IIΔ^UBC-ERT2)^ to induce the widespread deletion of MHC-II at late time points after infection. MHC-IIΔ^UBC-ERT2^ and control mice were infected with influenza followed by tamoxifen administration beginning at least 40 days after infection (Fig. 7d). Seven days after the last tamoxifen treatment, both resident and circulating B cells isolated from the lungs of MHC-IIΔ^UBC-ERT2^ mice expressed approximately 10-fold lower levels of MHC-II compared to control antigen presenting cells (Supplementary Fig. 7a, b). Although the number of B cells in the lung was not impacted by MHC-II deletion, the proportion and number of TRH cells was significantly decreased (Fig. 7e-g, Supplementary Fig. 7c, d). These data indicate that while both TRH and TRM1 cells are actively proliferating to a similar extent at late time points after infection, TRH are uniquely dependent on sustained antigen presentation for their maintenance.

**Fig. 7.**
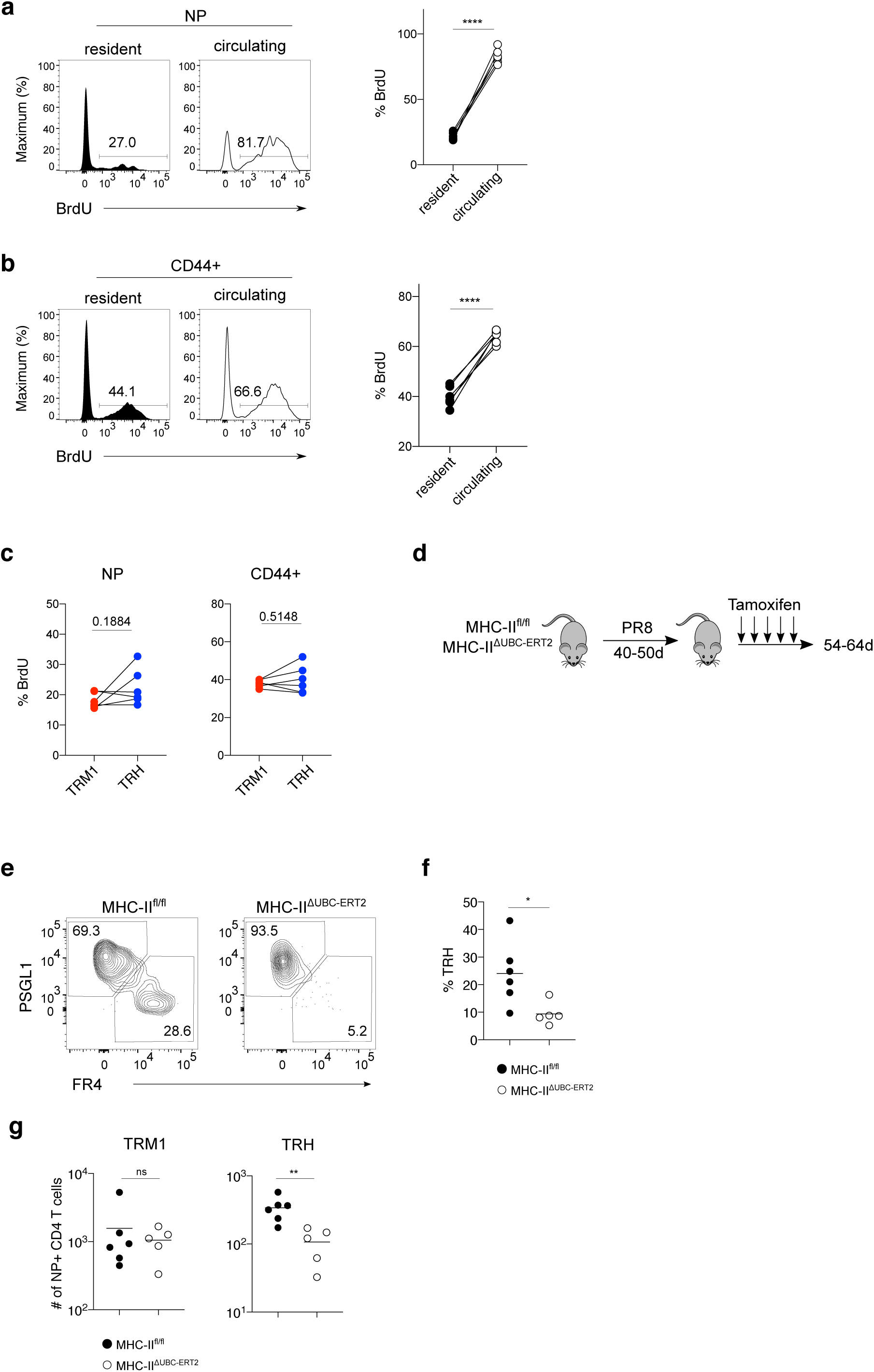
Maintenance of TRH cells requires antigen presentation. **a, b** Histograms and frequency of BrdU^+^ cells in NP-specific **a** and CD44^+^ **b** lung resident and circulating memory cells. **c,** Frequency of BrdU+ cells in TRM1 and TRH. **d,** Infected MHC-II^fl/fl^ and MHC-II^ΔUBC-ERT2^ mice were analyzed one week after inducible deletion. **e-f,** Flow cytometry plots **e** and TRH frequency **f** in MHC-II^fl/fl^ and MHC-II^ΔUBC-ERT2^ mice. **g,** Total numbers of NP-specific TRM1 and TRH. Data in **a**, **b** and **c** were analyzed by paired t-tests (n=5, representing 3 experiments). Data in **f** and **g** were analyzed by unpaired Student’s t-test (n=6, representing 2 experiments). Thin lines represent mean. P values are as follows: *P<0.05, **P<0.01, ***P<0.001, ****P<0.0001.

### TRH cells contribute to protection during re-challenge

TFH memory cells express high levels of *Tcf7*, a transcription factor associated with self-renewal and stem-like properties, and were recently shown to generate multiple cell fates following recall infection^26^. The similarities between TRH cells and TFH cells prompted us to investigate lung resident T cells for potential developmental trajectories. We chose TRH as a starting cluster for the analysis because it most closely resembles stem-like, *Tcf7*^hi^*Id3* ^hi^ TFH cells. Using PHATE for dimension reduction, we observed a bifurcating trajectory starting with the TRH cluster, branching at the *Hif1a* ^hi^ cluster 1 and terminating in the “circulating” cluster 4 and the TRM1 cluster 2 (Fig. 8a, Supplementary Fig. 8a). Imputed expression of genes emblematic for each terminal cluster could then be followed along developmental pseudotime (Fig. 8b). While we cannot exclude the possibility of circulating cells entering the lung as a starting point for a trajectory, enrichment of *S1pr1* and *Ccr7* in cluster 4 could also mark a “draining” population of cells that are poised to exit to the LN. These data raise the possibility that TRH, like TFH, may be multipotent following recall infection. To test this experimentally, we adoptively transferred TRH and TRM1 cells from influenza infected mice into congenic recipients followed by recall infection with PR8 influenza. At day 9 after challenge infection, both transferred TRM1 and TRH cells maintained high expression of PSGL1, similar to endogenous NP-specific cells at this time point (Fig. 8c, Supplementary Fig. 8a, b).

**Fig. 8.**
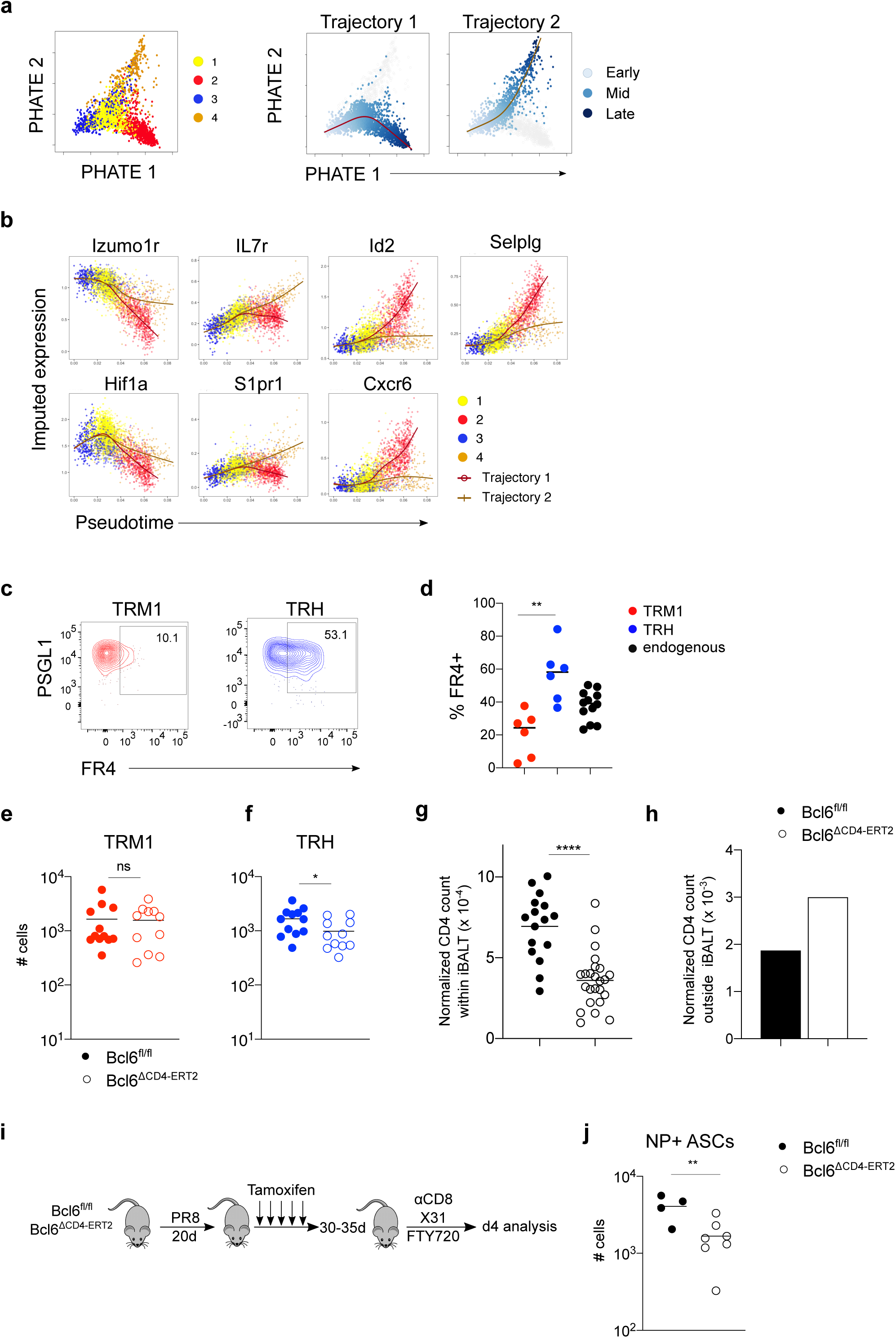
TRH cells contribute to protection during re-challenge. **a,** PHATE dimension reduction used as input to slingshot trajectory inference. PHATE colored by cluster (left) and pseudotime (right). **b,** Imputed expression of selected genes across pseudotime and two trajectories. **c,** Representative plots of FR4^+^ cells in TRM1 and TRH recipients. **d,** Frequencies from **c**. **e,** Total numbers of NP-specific TRM1. **f,** Total numbers of NP-specific TRH**. g,** Count of CD4 T cell objects inside B220+ iBALT clusters, normalized by iBALT volume, analyzed by Mann-Whitney-Wilcoxon test. Imaris segmentation. Each dot represents one iBALT. **h,** Count of iBALT-external CD4 T cell objects across all images, normalized by total tissue volume less iBALT volume. n = 2. **i,** PR8-infected, tamoxifen-treated Bcl6^flox/flox^ and Bcl6Δ^CD4-ERT2^ memory mice were treated with FTY720 and α-CD8 prior to X31 re-challenge and analyzed 4 days later. **j,** Number of NP-specific IgG ASC per mouse lung. Data in **c** (n=6, pooled from 2 experiments), **e** (n=11-12, pooled from 3 experiments) and **h** (n=4-7, representing 2 experiments) were analyzed by unpaired Student’s t-test. Thin lines represent mean. P values are as follows: *P<0.05, **P<0.01, ***P<0.001, ****P<0.0001.

Although the majority of transferred TRM1 cells remained negative for FR4 expression, transferred TRH cells gave rise to both FR4^hi^ and FR4^lo^ secondary effectors (Fig. 8c, d). These data demonstrate that similar to TFH memory cells, TRH cells retain the capacity to generate both FR4^hi^ and FR4^lo^ effectors while TRM1 cells appear to be more terminally differentiated at this time point.

TFH memory cells are reported to provide accelerated help to memory B cells during a challenge infection, in a T cell-intrinsic, Bcl6 dependent manner^39, 40^. As influenza infection can induce the differentiation of resident memory B cells in the lung, we next assessed the contribution of TRH cells to secondary humoral immunity in the tissue^41^. To do this we generated Bcl6^flox/flox^ x CD4^Cre-ERT2^ (Bcl6Δ^CD4-ERT2^) mice that allow for tamoxifen inducible deletion of Bcl6 in CD4 T cells. Bcl6Δ^CD4-ERT2^ and Bcl6^flox/flox^ control mice were infected with influenza and treated with tamoxifen starting at 21 days. Prior to secondary infection, late deletion of Bcl6 in CD4 T cells led to a mild decrease in the number of NP-specific TRH cells while TRM1 cells numbers were not affected (Fig. 8e, f). To understand if Bcl6 deletion had an impact on CD4 resident T cell maintenance within iBALT, we examined the lungs of Bcl6Δ^CD4-ERT2^ and control mice by histology.

While CD4 cells were abundant in dense areas of strong B220 signal in control mice, Bcl6 deletion led to significantly reduced colocalization of these cell types, despite the maintenance of large clusters of B220+ cells (Fig. 8g, Supplementary Fig. 8b). In contrast, normalized CD4 cell counts outside of iBALT areas were higher with Bcl6 deletion (Fig. 8h). As iBALT can act as a site for the local priming of pulmonary immune responses we next examined whether Bcl6 deletion in CD4 T cells would lead to less efficient secondary B cell responses. To test this, we further treated Bcl6Δ^CD4-ERT2^ and control mice with FTY720 and a CD8 depleting antibody followed by challenge infection with heterotypic X31 influenza (X31) (Fig. 8i). This approach allowed us to examine the impact of Bcl6 deletion in lung CD4 T cells independently from circulating lymphoid T and B cells, as well as memory CD8 T cells, which might otherwise lead to early viral clearance. Importantly, antibodies generated after PR8 influenza infection do not neutralize X31 infection, while CD4 T cells respond to the conserved NP epitope, thereby enabling specific investigation of CD4 T cell mediated protection. Four days after recall with X31 influenza, Bcl6 deletion in the CD4 compartment led to a four-fold decrease in NP-specific antibody-secreting cells in the lung (Fig. 8j, Supplementary Fig. 8c). These data indicate that TRH cells contribute to heterologous protection by promoting local antibody production during re-challenge.

## Discussion

In this study we used scRNAseq to characterize the dynamics, heterogeneity and transcriptional regulation of lung resident CD4 T cells after influenza infection. In contrast to previous studies which mainly identified Th1 memory cells with transcriptional resemblance to CD8 TRM cells, our data reveal the presence of long-lived TRH cells with phenotypic and functional similarities to lymphoid TFH cells. In agreement with these earlier studies, many of which tracked monoclonal T cell responses with specificity to a single antigen, we did not observe TRH differentiation by OT-II T cells responding to PR8-OVA, indicating that not all TCR lines recapitulate the heterogeneity of polyclonal T cell populations. Importantly, our study identifies phenotypically similar cell clusters in both LN and lung alongside a consistent signature of residency in the lung. Given the predominance of type 1 responses to viral infection in barrier tissues, the averaging effect of bulk studies, and the paucity of antigen-specific polyclonal models to date, a general residency signature was previously obscured.

Notably, the TRH population we identified in the lung was itself heterogeneous, with FR4^hi^PSGL1^low^ cells clustering into two groups transcriptionally. Regulation of these two subsets by Bcl6 and HIF-1α, respectively, is consistent with a previous study showing that Bcl6 can repress gene programs activated by HIF-1α in effector CD4 T cells^42^. While HIF-1α has been alternately reported to repress or promote TFH effector cell differentiation^43, 44^, our data reveal a gradual upregulation of HIF-1α in CD4 TRM cells, coincident with TRH cell differentiation, localization in iBALT and resolution of the inflammatory/hypoxic phase of infection. HIF-1α expression might therefore reflect differences in TRH cell localization and the dynamics of access to environmental signals. In addition, expression of *Areg*, encoding the cytokine amphiregulin that promotes epithelial cell proliferation, may also point to a role in tissue homeostasis^45^.

Amphiregulin produced by regulatory T cells was previously shown to prevent excessive inflammation in the lung at very early time points after primary infection^46^. It will be interesting to determine if amphiregulin produced by resident CD4 T cells plays a similar role during recall infection, as a means to balance anti-pathogen immunity with tissue integrity and niche preservation.

Our data highlight the importance of B cell interactions for the generation of the TRH compartment. B cells have been previously implicated in the differentiation of Th1 TRM cells following intranasal LCMV infection, where they initially restrained Th1 cell residency but later promoted their long-term persistence^47^. Although these results are consistent with our observation that TRM1 cells differentiate earlier compared to TRH cells, we did not observe any impact on TRM1 cell numbers in B cell depleted mice. We additionally determined that TRH cells require intrinsic Bcl6 expression. The absence of Bcl6 at the start of infection led to complete ablation of the TRH compartment as well as a significant decrease in TRM1 cells. These data are consistent with a previous study looking at CD4 T cell differentiation in the lung during tuberculosis infection^48^. Here the authors reported that both Bcl6 and ICOS signaling are required for the development of lung resident T cells with memory like properties that mediate superior protection compared to more terminally differentiated lung Th1 cells. Taken together with our observations, these findings suggest that TRH cells maintain the plasticity to differentiate into Th1 effectors following challenge infection, an idea that is supported by our trajectory analysis and adoptive transfer experiments. The ability of TRH cells to differentiate into multipotent effectors is similar to stem-like TFH memory cells which were recently reported to generate diverse effectors following secondary infection^26^.

Influenza infection results in the development of iBALT, a tertiary lymphoid structure (TLS) in the lung which can act as an immunological hub, promoting rapid and localized immune responses following secondary infection^49^. Compared to TRM1 cells, we detected tight localization of TRH cells and B cells within iBALT cores. The importance of T-B interactions in TLS has been highlighted in many immune contexts, including tuberculosis infection, where the presence of iBALT correlates with bacterial control, as well as the tumor microenvironment, where the presence of TLS predicts improved patient outcome^50–54^. Our data provide additional insight to these studies, demonstrating TRH cell dependency on both antigen presentation and Bcl6 expression for their maintenance within iBALT. The decreased number of iBALT-localized CD4 T cells observed after late Bcl6 deletion may be due to lower expression of ICOS which normally promotes T cell interactions with ICOSL expressing B cells^55^. Further experiments will help clarify if Bcl6 is also required for the long-term maintenance or retention of CD4 memory cells in other TLS contexts such as the tumor microenvironment or autoimmune diseases such as rheumatoid arthritis, where TFH cells are implicated in health and pathology, respectively.

Finally, our data demonstrate an important role for resident CD4 T cells in orchestrating local humoral responses during recall infection. This is reciprocal to the relationship between memory TFH cells and memory B cells present in lymphoid organs where antigen specific memory B cells induce Bcl6 expression in cognate TFH cells, leading to more efficient T cell help and acceleration of humoral immune responses^39, 40^. Our experiments suggest that TRH cells are poised to provide help due to their tight localization with BRM cells. Thus, the targeted induction of lung TRH cells may serve as the basis for rational design of a universal influenza vaccine. Taken together with the observation that TRH cells maintain the capacity to generate Th1 effectors, these findings have implications for reinvigorating or dampening T cell responses in tissues or tumors where TLS are present.

## Methods

### Mice

Male and female mice C57BL/6 J (CD45.2), B6.SJL-Ptprc^a^Pepc^b^/BoyJ (CD45.1), C57BL/6-Tg(Nr4a1-EGFP/cre)820Khog (Nur77 GFP), Bcl6^tm1.1Cdon^ (Bcl6 RFP), Tg(Tbx21-ZsGreen)E3ZJfz (T-bet ZsGreen), B6.129X1-H2-Ab1^tm1Koni^/J B6.Cg-Ndor1^Tg(UBC-Cre/ERT2)1Ejb^/1J (MHCIIΔ^UBC-ERT2^), (UBC-ERT2 mice kindly provided by Prof. Tobias Derfuss, University of Basel), B6.Bcl6^tm1.1Mtto^Tg(Cd4-cre)1Cwi (Bcl6Δ^CD4^), B6.Bcl6 ^tm1.1Mtto^ Cd4^tm1(cre/ERT2)Thbu^ (Bcl6Δ^CD4-ERT2^, Bcl6 fl/fl mice kindly provided by Dr. Toshitada Takemori, RIKEN The Institute of Physical and Chemical Research), B6.Cg-Tg(TcraTcrb)425Cbn/J (OT2) were used. Mice in each experiment were same sex littermates, 6 to 12 weeks old at the start of experiment, maintained and bred in the specific pathogen free animal facility at the University of Basel. All animal experiments were performed in accordance with local and Swiss federal guidelines. Non-blinded experimental groups were formed with random assignment of mice and no specific methods applied to determine sample size.

### Infections

Influenza A/PR8/34 OVAII (hereafter, PR8) and X31 starter material were kindly provided by Dr. Paul G. Thomas, St. Jude Children’s Research Hospital and produced at VIRAPUR. PR8 was intranasally administered at 500-2000 TCID_50_ for all primary infections. Influenza X31 was used at 10^5^ TCID_50_ for secondary infections. Mice were anesthetized with vaporized isoflurane and infected intranasally with virus diluted in 30-40μL volume of PBS.

### Mixed bone marrow chimera

Bone marrow cells from wild type (CD45.1) and Bcl6^ΔCD4^ (CD45.2) mice were collected, mixed in 60:40 (CD45.1:CD45.2) ratio to compensate for reconstitution defects of Ly5.1 line and adoptively transferred into lethally irradiated (2x 500cGy) CD45.1 hosts^56^. Reconstitution in blood was checked 6 weeks after irradiation.

### Adoptive transfer

Inguinal, brachial and axillary lymph nodes from naive OT2 mice were harvested and mashed through a 100μm Nylon cell strainer (Corning). Negative selection for CD8α, CD19, B220, I-A^b^, NK-1.1 using LS columns (Miltenyi Biotec) was performed to enrich CD4 T cells. Cells were enumerated as previously described and 30,000 OT2 cells were transferred into congenic hosts one day before infection with PR8-OVA2^57^. For the cell transfer experiment in Fig. 8c, non-circulating CD4 T cells from each subset were sorted and transferred into naive congenic hosts that were infected with PR8 the following day.

### In vivo treatments

Tamoxifen (Sigma) was dissolved in corn oil (25mg/ml) and stored at 4°C during the duration of treatment. Mice were administered 3.75mg in 150μl volume for 5-7 days by oral gavage. Mice were administered BrdU (10280879001 Roche) in the drinking water (1mg/ml with 1% glucose) for 13 days and sacrificed on the fourteenth day. 12.5μg homemade ARTC2.2-blocking nanobody s+16 (NICD protector) was intravenously injected into mice at least 15 min before sacrifice. Mice were injected intravenously with 3μg of fluorochrome conjugated anti-CD45 AF700 (clone A20, #110724, Biolegend) 3 min prior to sacrifice to label circulating cells and discriminate them from the non-circulating compartment. B cell depletion was performed by weekly i.p. administrations of 250μg of murine B cell depletion antibody (anti-mouse CD20, 18B12 obtained from Biogen MA Inc.) starting from one week before infection until sacrifice. CD8 depletion was performed by administering InVivoMAb anti-mouse CD8α (clone 2.43, BioXCell), 400μg i.p. and 100μg intranasal for 2 consecutive days before secondary infection. The S1P1R agonist, FTY720 (AdipoGen Life Sciences) was dissolved in DMSO at a concentration of 20mg/ml. 1mg/kg was administered i.p. for three consecutive days before harvest for memory time point experiments (Supp. Fig 1), every 2 days starting day 9 for migration experiment (Fig. 4) and every day starting 3 days before secondary infection for recall experiments.

### Tissue preparation

Lungs were harvested and diced into GentleMACS C Tubes (Miltenyi Biotec) and washed down with 3ml media (RPMI, 10mM aminoguanidine hydrochloride (Sigma), 10mM HEPES, Penicillin-Streptomycin-Glutamine (100X, Gibco), 2-Mercaptoethanol (50μM, Gibco)) devoid of FCS and EDTA pre-warmed in 37°C water bath. Digestion mix containing 33.3μg/ml Liberase (Roche #05401020001) and 68μg/ml DNAse I (Applichem) were added. Lungs were dissociated on gentleMACS Dissociator (Miltenyi Biotec) and placed in a shaking incubator at 37°C for 30 min. Lungs were dissociated again and mashed through 70μm MACS SmartStrainers (Miltenyi Biotec). Red blood cells were lysed with 139.5mM NH_4_Cl and 17mM Tris HCl (Erylysis buffer). For further processing, media containing 2% FCS and 1mM EDTA were used. Mouse mediastinal lymph nodes were harvested and mashed through 100μm Corning Nylon cell strainer.

### Tetramer, antibody staining, flow cytometry

I-A^b^ NP311-325 (QVYSLIRPNENPAHK) Allophycocyanin was obtained from the NIH tetramer core facility. Single cell suspensions were tetramer stained, enriched and counted as previously described^57^. Dasatinib (50nM) was added to the tetramer mix to reduce pMHC tetramer binding affinity threshold and activation induced cell death^58^. To analyze B cells, the flowthrough collected post tetramer enrichment was used to enumerate and stain for B cell markers. For all fluorochrome-conjugated antibody dilutions, FACS buffer (PBS, 2% FCS, 0.1% Sodium Azide) containing Fc block (InVivoMAb anti-mouse CD16/CD32, Clone 2.4G2, BioXCell) was used. Live-dead stain (Zombie Red Fixable Viability kit, Biolegend) was added to the antibody mix and stained at 4°C for 30 min. FoxP3 fixation kit (eBioscience) was used for intracellular staining at room temperature (RT) for 1 hour. For BrdU staining, BD Pharmingen BrdU Flow Kit was used. Flow cytometric analysis was performed on BD LSR Fortessa. Data were analyzed using FlowJo X software (TreeStar).

### Gating strategy used in Flow cytometric analysis

To gate on resident CD4 T cells, lung cells were gated on Zombie red^-^ (live), dump^-^ (CD11b, CD11c, B220, F4/80) lymphocytes and then on CD4^+^, i.v^-^ T cells (non-circulating). For antigen-specific T cells, CD44^+^NP^+^ cells were gated on. Non-antigen specific are considered as CD44^+^NP^-^. The same gating strategy was used for CD4 T cells from mLN.

For B cell staining in the lung, cells were gated on Zombie red^-^ (live), dump^-^ (CD3ε, Gr-1, F4/80, CD11c) lymphocytes and then on B220^+^. Further gating on iv^+^ or iv^-^ was done to discriminate between circulating and non-circulating B cells.

### scRNA sequencing

0.5-3 x 10^4^ total NP-specific CD4 T cells were sorted from mLN and lung of PR8-OVA2 infected mice at indicated time points and were provided for library preparation using the 10x Chromium platform. Each sample is pooled from 4-12 C57BL/6 J mice. Sorting was performed using BDFACSAria^TM^ III and BDSorpAria^TM^ III. Single-cell capture and cDNA library preparation were performed with a Single Cell 3’ v2 Reagent Kit (10x Genomics) according to manufacturer’s instructions. Sequencing was performed on one flow-cell of an Illumina NexSeq 500 at the Genomics Facility Basel of the ETH Zurich. Paired-end reads were obtained and their quality was assessed with the FastQC tool (version 0.11.5). The length of the first read was 26 nt, composed of individual cells barcodes of 16 nt, and unique molecular identifiers (UMIs) of 10 nt. The length of the second read, composed of the transcript sequence, was 58 nt. The samples in the different wells were identified using sample barcodes of 8 nt.

### scRNA sequencing analysis

Sequencing data was processed using 10X Genomics’ cellranger software version 2.1.0, modified to report only one alignment (randomly) for multi-mapped reads (keep only those mapping up to 10 genomic locations). Raw molecule info from cellranger was initially filtered leniently, discarding cells with fewer than 100 UMI counts, as well as the highest 99.99% to account for likely doublets. The resulting UMI matrix was further filtered to keep only cells with log library size > 2.9, log number of features > 2.6, percent mitochondrial reads <= 6, and percent ribosomal protein reads >= 20. Genes with average counts < .007 were removed. Normalization was done using the R package scran’s deconvolution method. Technical noise within gene expression was modeled using scran, and biologically relevant highly variable genes were calculated after separating the technical from biological variance, using FDR < 0.05 and biological variance > 0.1. PCA was run on the normalized data using the top 500 most variable genes by biological variance, and the PCA was denoised to account for the modelled technical variation. Cells were clustered hierarchically using Ward’s method on the distance matrix of the PCA. Default dendrogram cut height using the R package dynamicTreeCut resulted in clusterings that also mapped to the highest average silhouette width. Data subsets, e.g. lung d30 cells only, were subjected to the same pipeline after subsetting. The sex-linked gene *Xist* was removed from the signature residency gene list. Subsequent visualization and analysis were performed using version 3.1.1 of the Seurat R package; dropout imputation was performed using Seurat version 2.3.4. Single-cell regulatory network inference and clustering (SCENIC) was performed using version 0.9.7 and the published workflow on the pySCENIC github repository (96). Pseudotime analysis was performed with slingshot version 1.5.0 and PHATE dimension reduction^29, 59^. Secondary analysis of published data sets was taken directly from published supplemental material or downloaded from GEO and analyzed for differential expression between samples by the limma package^60^.

### Histology

Lungs were inflated with 3% low melting agarose (LMA) (Sigma, #A9414), fixed in 1% formaldehyde (Thermo scientific, #28908) for 24h at 4°C, washed with PBS and kept at 4°C in PBS + 0.01%NaN_3_. Lung tissues were embedded in 3% LMA and 100μm-thick sections were cut at the vibratome (Leica VT 1200S). Sections were placed on microscope slides, circumscribed with ImmEdge pen (Vector) and blocked in TBST (100mL Tris-HCl, 150mM NaCl, 0.05% Tween-20) containing 5% Donkey serum (Jackson ImmunoResearch #017-000-121) for 2 hours at RT. For lung staining from T-bet ZsGreen mice, sections were incubated with anti-CD4 (1:50 in blocking buffer) overnight at 4°C, then washed (4 times for 15 min with TBST) and incubated with anti-rat-AF647 (1:160) for 4 hours at RT, followed by washes and subsequent incubation with anti-B220-BV421 (1:50) overnight at 4°C. For Bcl6 RFP mice, sections were processed similary by incubating sections with anti-CD4 (1:50) and anti-RFP (1:100) overnight at 4°C, then with anti-rat-AF647 (1:160) and anti-rabbit-AF488 (1:200) for 4 hours at RT, and finally with anti-B220-BV421 (1:50) overnight at 4°C. For Bcl6^fl/fl^ and Bcl6Δ^CD4-ERT2^ mice, sections were incubated with anti-CD4 (1:50) overnight at 4°C, then washed, incubated with anti-rat-AF647 (1:160) for 4 hours at RT. Then sections were washed, incubated for 45 min with 5% rat serum in blocking buffer, washed, incubated for 45 min with Fab anti-rat (100μg/ml), washed and incubated with anti-B220 (1:50) overnight at 4°C. Sections were then washed and incubated with anti-rat-AF488 (1:300) for 4 hours, RT. After the last wash, sections were mounted using ProLong Glass antifade mountant (Invitrogen, P36984).

### Microscopy

Images (16bts, 1000×1000 pixels images) were acquired using X20 or X40 magnification objectives at the spinning disc confocal microscope (Nikon CSU-W1) with a photometrics 95B camera.

### Image quantification

Confocal images were processed in Bitplane Imaris v9.5. Surfaces were created to segment different features of interest: dense B cell areas (B220+) (automatic threshold, surface detail 8 µm, seed point diameter >= 40, artefacts removed and final areas unified manually), individual CD4+ T cells (automatic threshold, surface detail 0.8 µm, seed point diameter 4 µm), and the total possible volume in which a CD4+ T cell could be found (threshold ∼150, surface detail 8 µm). Where necessary, surfaces for obvious non-cell artefacts were created and used to mask the channels of interest before creating target surfaces. The statistics including shortest distance between surfaces was exported to CSV files. To confirm these results, images were also processed using custom computer vision and machine learning algorithms mainly written in python. Two separate image-based classifiers were trained at voxel and object level detection and identification for 3D clusters (B220) and 3D single cells (CD4). Features such as intensity, morphology, etc., were then measured per object at cluster and cellular level and colocalized using a parent (cluster) and child (cell) relationship within clusters of B cells. The distances between surfaces were calculated using centroids of cells and clusters. Data from both segmentation pipelines were loaded into R and processed using custom code to quantify and test distributions of CD4 T cells with respect to B cell areas. CD4 objects less than approximately 2.5 µm in radius were discarded as probable artefacts of surface creation. B cell clusters smaller than 20000 µm^3^ were also discarded. Double positive CD4+Tbet+ or CD4+Bcl6+ cells were positively gated as if for FACS based on a log distribution of mean intensity scores per CD4 object - by eye with the manual pipeline and at a threshold of the top 10% of per image data for the machine learning pipeline. Normalization: T cell counts within B cell areas were divided by that B cell area’s volume, and counts outside B cell areas were divided by the total available T cell volume minus all B cell areas.

### ELISpot

Multiscreen IP HTS filter plates (#MAIPS4W10, EMD Millipore) were coated with 2μg/mL NP (Sino biological #11675-V08B) and incubated overnight at 4°C. Plates were blocked with cell culture medium. Cells were plated and incubated at 37°C for 5 hours. Plates were washed in PBS+0.01% Tween-20 before goat anti-mouse IgG HRP (Fcγ fragment, Jackson Immuno #115-035-008) was added and incubated overnight at 4°C. Plates were washed and developed using AEC Substrate set (BDBiosciences #551951). Spots were recorded with AID Elispot Reader and enumerated manually.

### Statistical analyses

Statistical analyses were performed with Prism Graphpad Software (Version 8.0). Unpaired or paired t-tests (two-tailed) or Mann-Whitney-Wilcoxon tests were used according to the type of experiments. P < 0.05 was considered significant.

## Acknowledgements

We thank D. Pinschewer and his lab for helpful discussions; J. Roux and F. Geier for expertise and technical advice; R. Tussiwand and her lab for feedback and support; the flow sorting facility and all the animal caretakers at the DBM University of Basel. Single cell RNA sequencing was performed at the Genomics Facility Basel, ETH Zurich. Calculations were performed at sciCORE (http://scicore.unibas.ch/) scientific computing center at the University of Basel.

## Data and materials availability

scRNAseq data are deposited with the National Center for Biotechnology Information Gene Expression Omnibus (accession nos: PLACEHOLDER).

---

**Supplementary Fig. 1.**
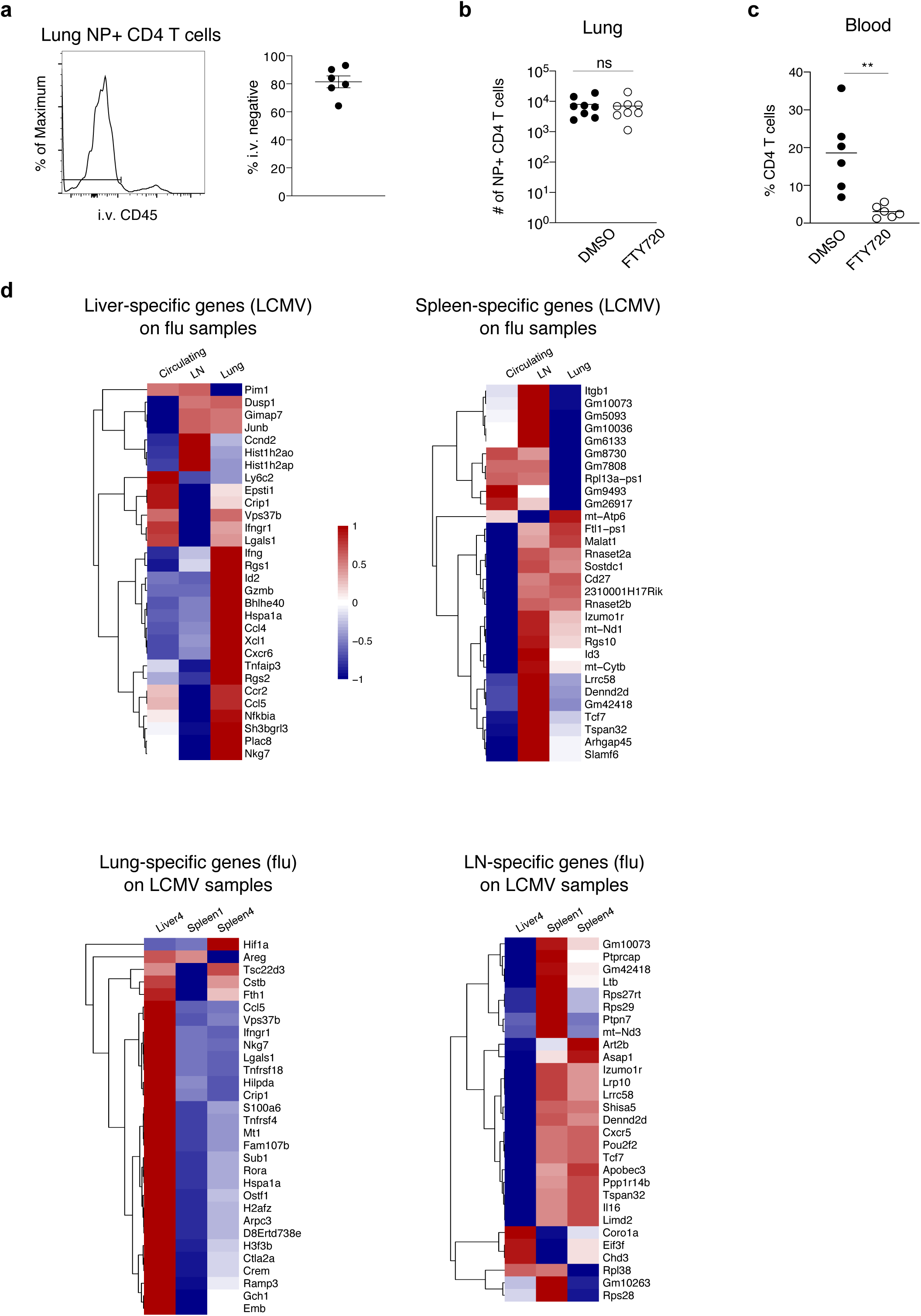
Residency signature in CD4 T cells is biased toward Th1 memory cells. **a,** Identification and frequency of NP-specific resident T cells. **b-c,** Total numbers of NP-specific T cells (**b)** and total CD4 T cell frequencies in blood (**c)** of control and FTY720-treated mice. **d,** Tissue specific signatures derived from one infection model plotted on data set from the other infection model. Heatmaps show cluster-averaged, scaled centered expression.

**Supplementary Fig. 2.**
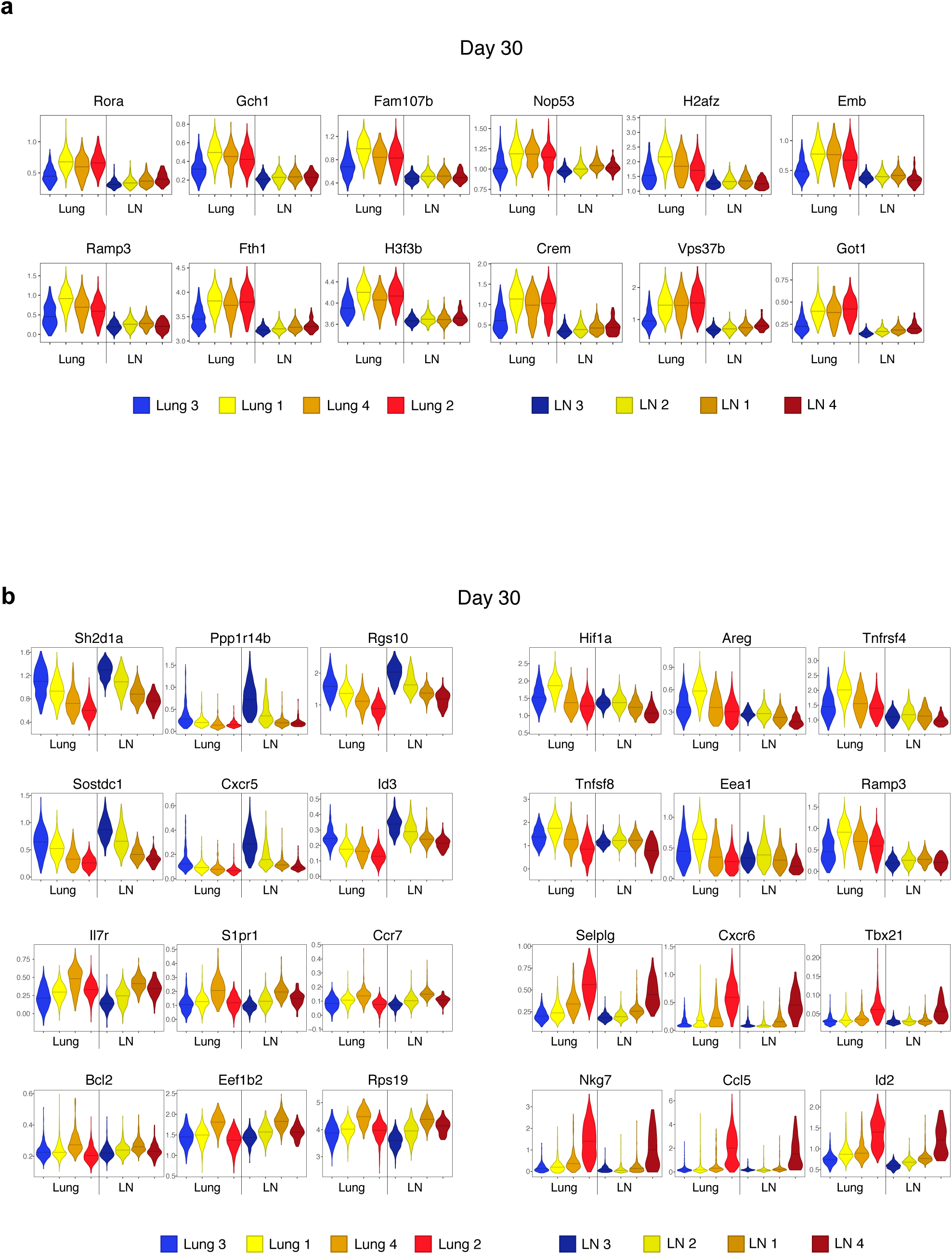
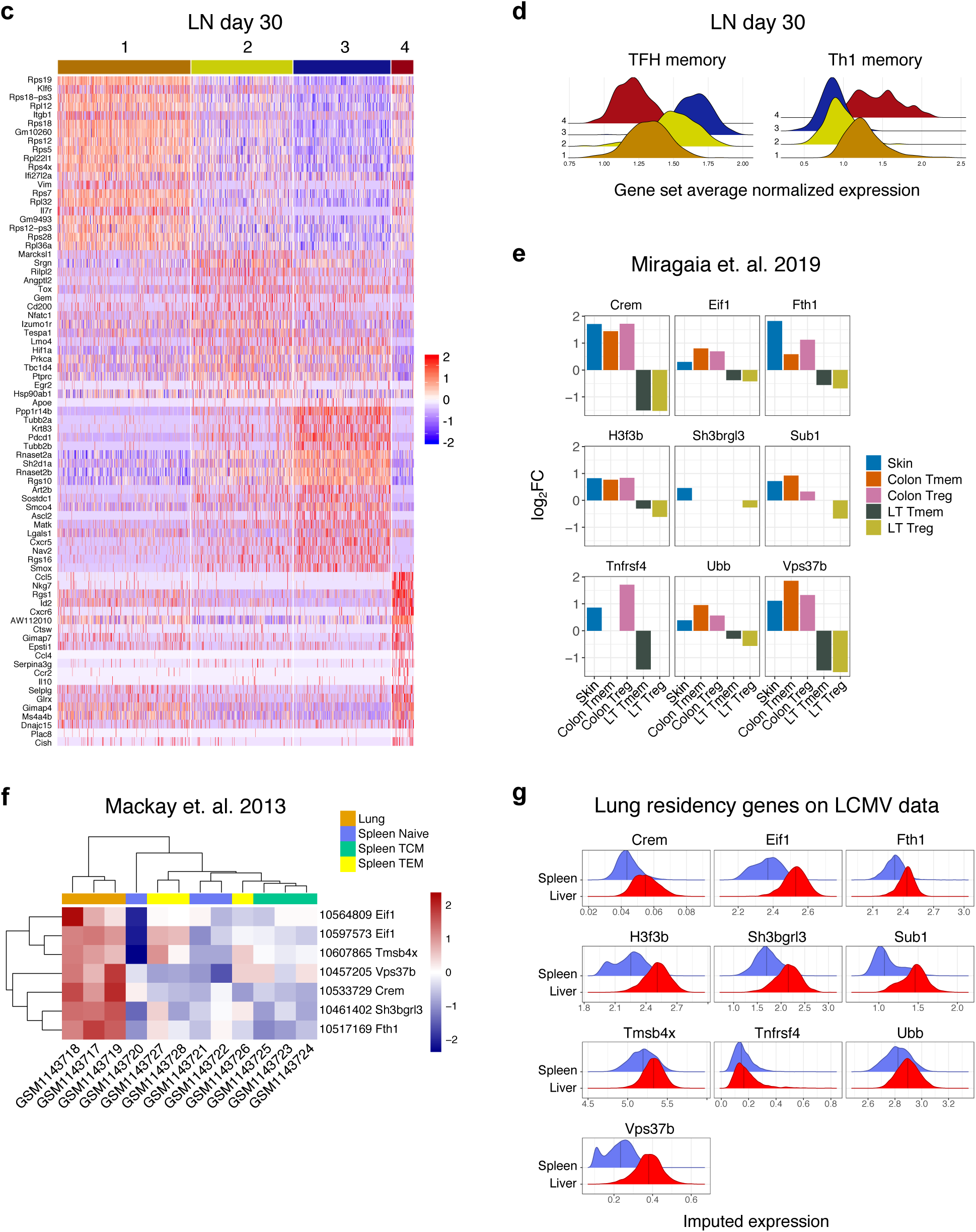
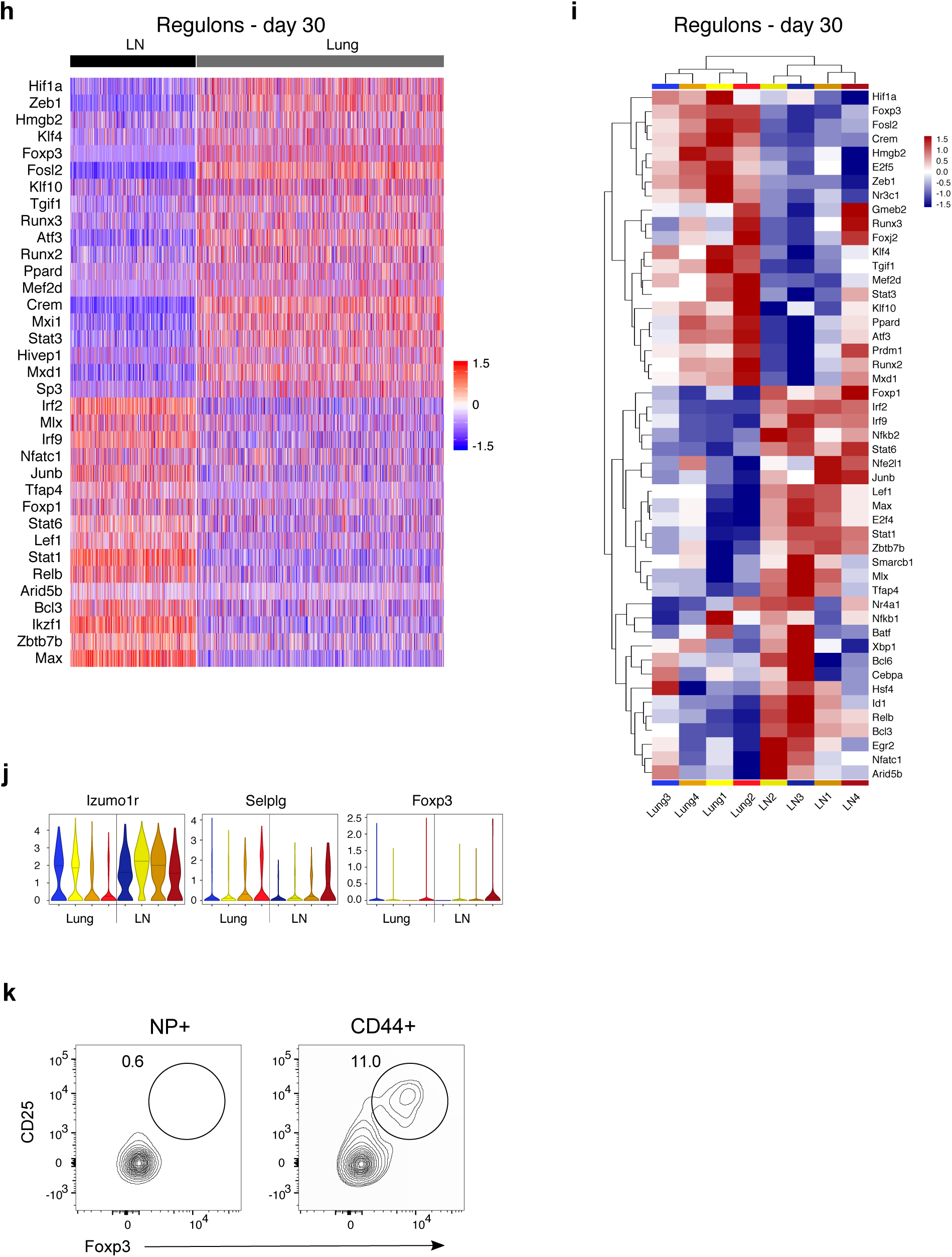
Heterogeneity in the lung: pan residency versus Th-subset specific residency. **a,** Imputed expression of residency genes. Dropout imputed using Seurat::AddImputedScore. **b,** Imputed expression of genes typifying similar clusters in lung and LN: blues = TFH-like, yellows = TFH-like, oranges = TCM-like, red = Th1-like. **c,** Heatmap showing scaled, centered single cell expression of top 20 genes sorted according to LN cluster average log_2_FC, adjusted P value < 0.05. **d,** Log-normalized average expression of TFH and Th1 memory signatures^26^. **e,** Conserved tissue residency signature genes on published single cell data set analyzing Treg adaptation to tissue^11^. **f,** Conserved tissue residency signature genes on published data set investigating CD8 TRM^62^. **g,** Imputed expression of conserved tissue residency signature genes on LCMV data. **h,** Scaled, centered area under curve (AUC) calculated with SCENIC showing top 20 differentially active transcription factors by tissue at day 30. Adjusted P-value < 0.01. **i,** Scaled, centered, cluster average SCENIC AUC showing top 20 differentially active TFs by tissue and cluster. Adjusted P value < 0.01. **j,** Log-normalized expression by cluster. **k,** Foxp3+ cells in the lung. Data in **a** (n=6, representing 3 experiments), **b** (n=8, pooled from 2 experiments) and **c** (n=6, pooled from 2 experiments) were analyzed by unpaired Student’s t-test. Thin lines represent mean ± s.e.m in **a** and mean in **b, c**. Data in **i** represents 2 experiments, n=5. P values are as follows: *P<0.05, **P<0.01, ***P<0.001, ****P<0.0001.

**Supplementary Fig. 3.**
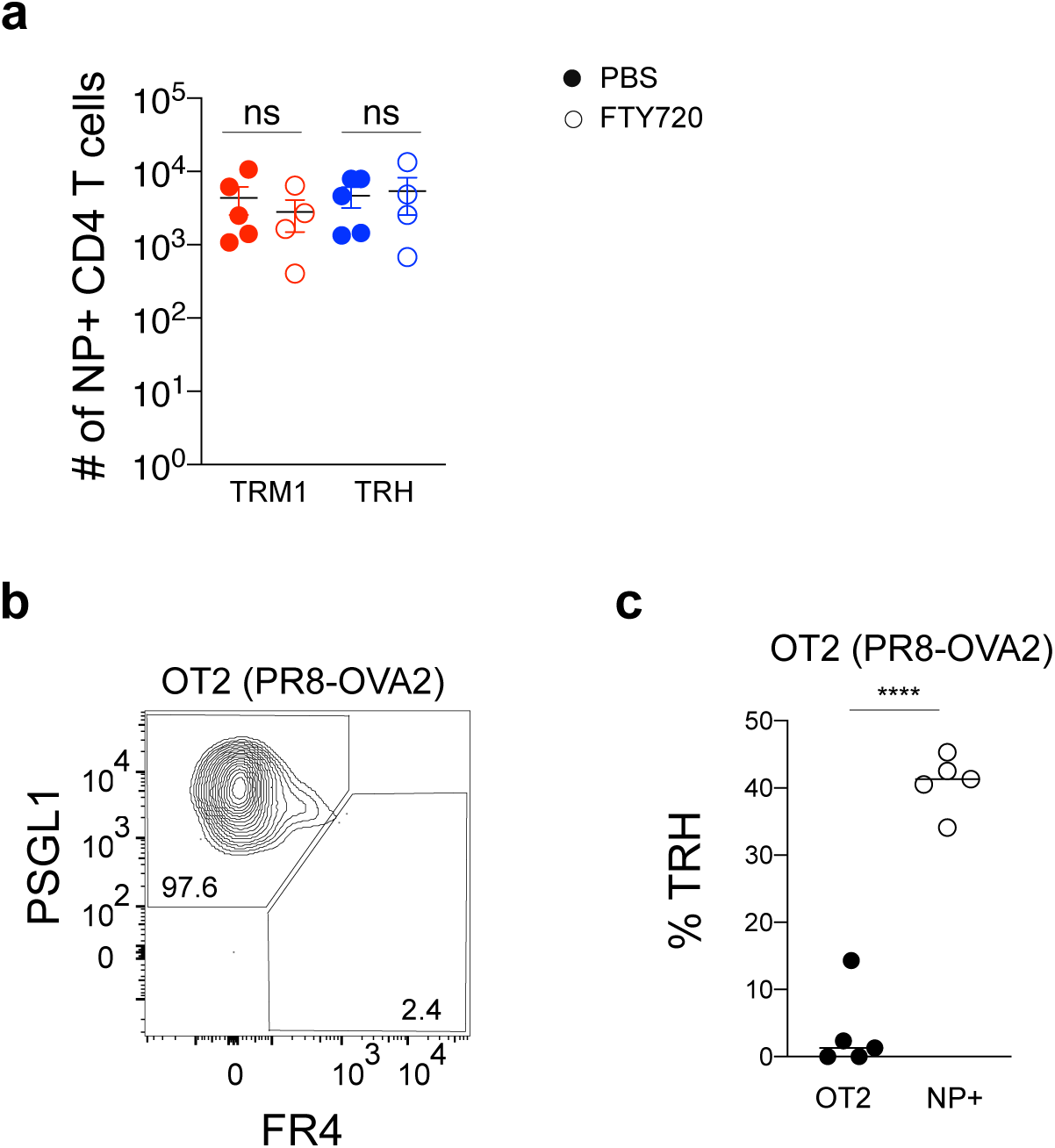
Phenotypic characterization of resident CD4 T cell subsets. **a**, Total numbers of NP-specific non-circulating TRM1 and TRH from PBS and FTY720-treated mice. **b, c** Representative plot **b** of TRM1 and TRH among transferred OT2 and frequency **c** of TRH in OT2 and endogenous NP-specific CD4 T cells. Data in **a** (n=4-5, representing 2 experiments) and **c** (n=5, representing 2 experiments) were analyzed using unpaired Student’s t-test. Thin lines represent mean± s.e.m in **a** and mean in **c**. Significance was determined using unpaired Student’s t-test. P values are as follows: *P<0.05, **P<0.01, ***P<0.001, ****P<0.0001.

**Supplementary Fig. 4.**
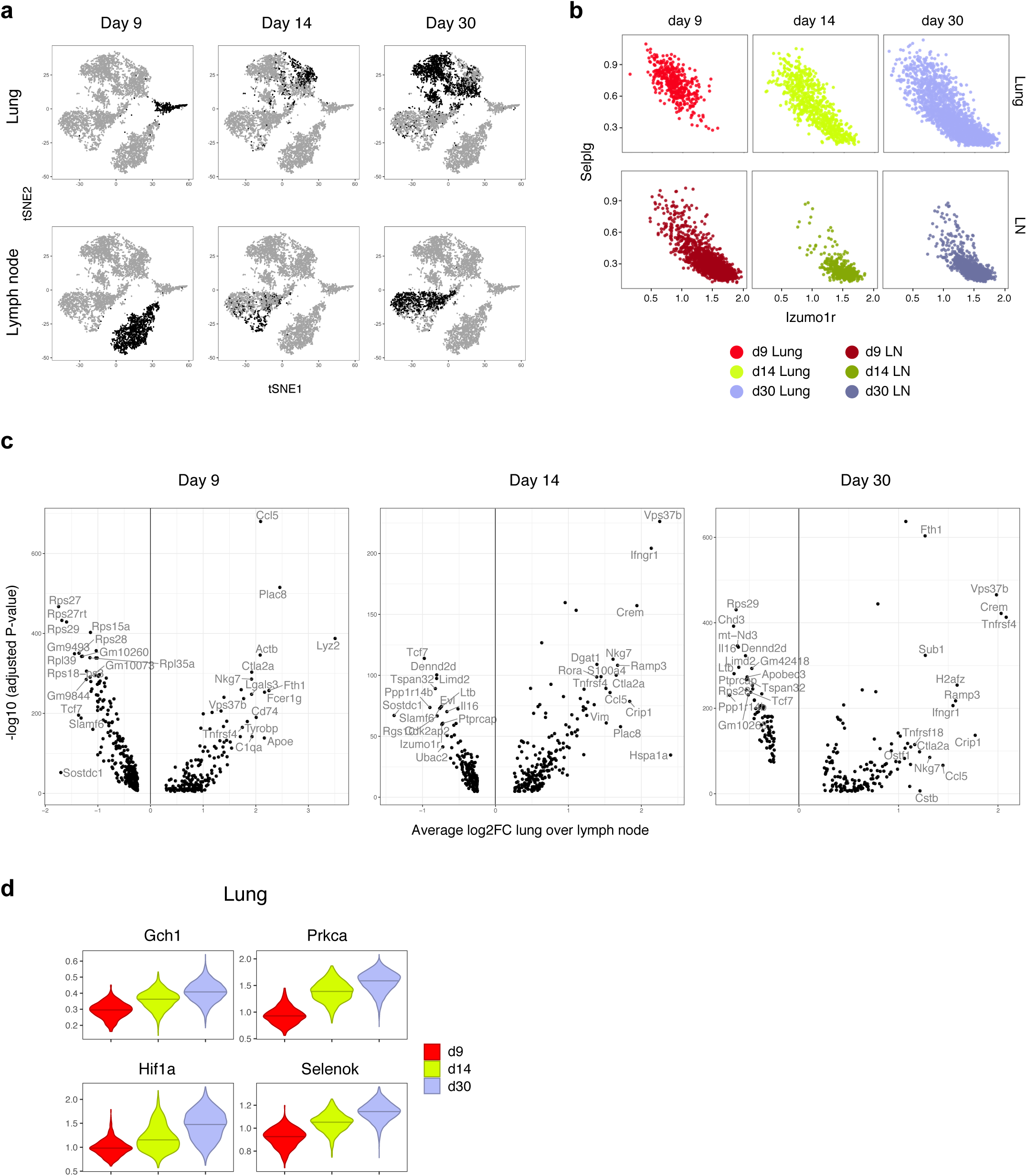
Progressive differentiation of resident CD4 T cell subsets. **a**, tSNE of scRNA-seq data by tissue and time point. **b,** Imputed expression of *Izumo1r* and *Selplg* by tissue and time point. **c,** Differentially expressed genes discriminating lung (+) from LN (-) across time points. **d,** Imputed expression of selected genes in the lung across time points.

**Supplementary Fig. 5.**
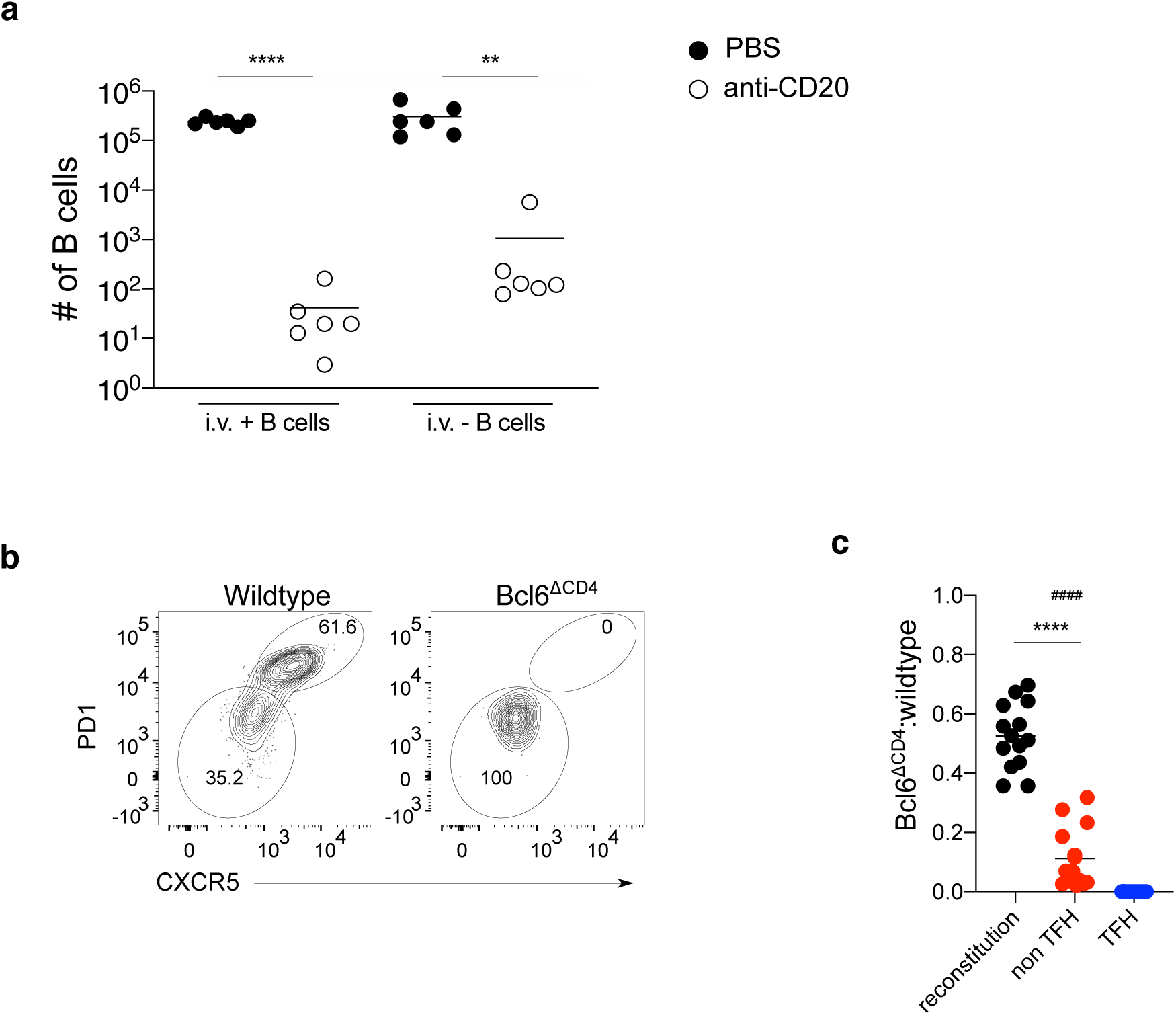
TRH cell generation requires B cells and T cell intrinsic Bcl6. **a,** Total numbers of iv^+^ and iv^-^ B cells in control and B cell depleted mice. **b,** NP-specific TFH and non-TFH cells of mLN from control and Bcl6-deficient subsets. **c,** Bcl6Δ^CD4^:control ratio of CD4 T cells after reconstitution from blood (pre-infection), NP-specific non-TFH and TFH. Thin lines represent mean in **(a)** (n=6, representing 2 experiments) and **(c)** (n=15, pooled from 3 experiments). Significance was determined by unpaired Student’s t-test. P values are as follows: *P<0.05, **P<0.01, ***P<0.001, ****P<0.0001.

**Supplementary Fig. 6.**
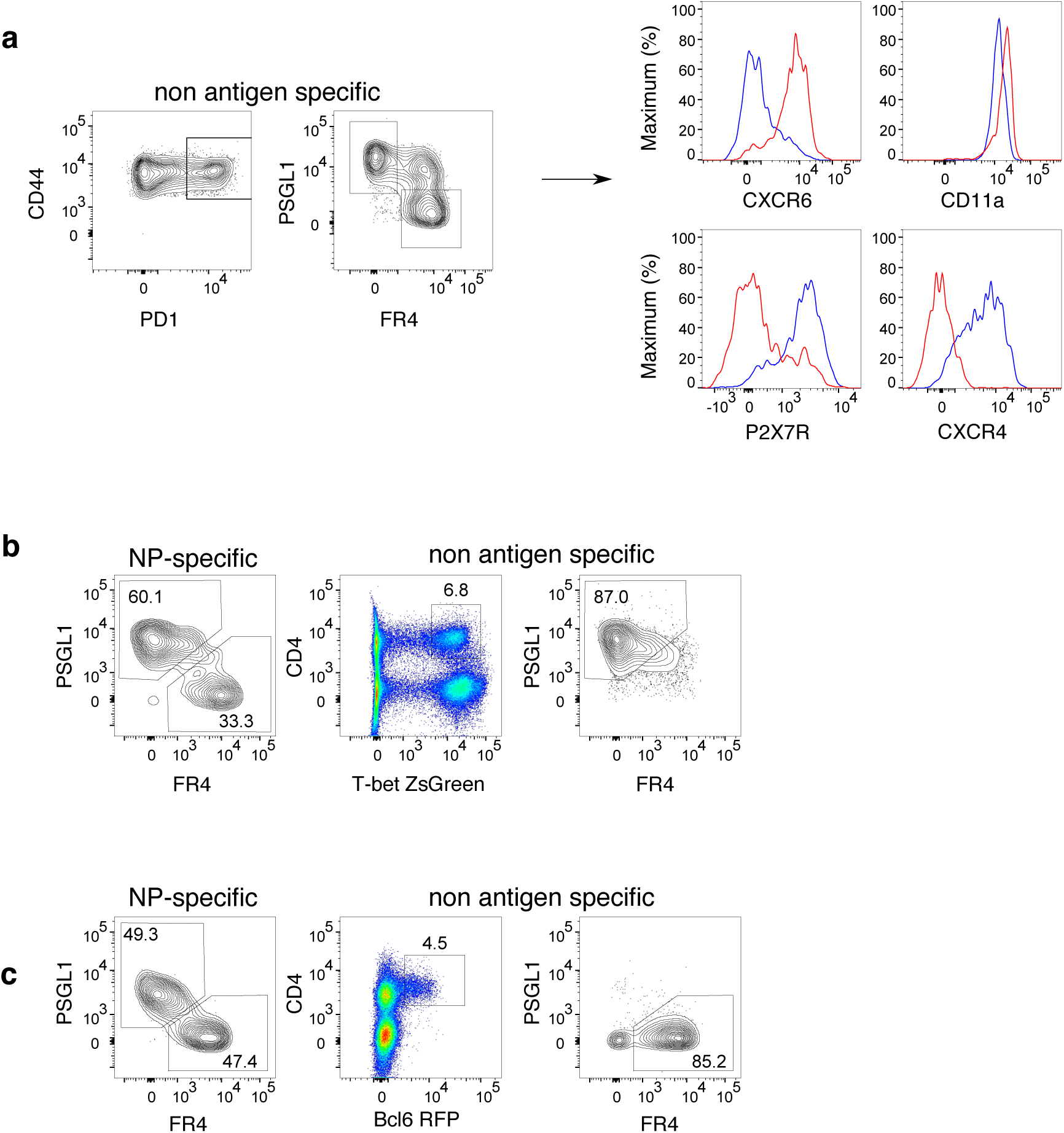
CD4 TRH cells localize in iBALT. **a,** Representative gating strategy for identification of non-antigen specific resident CD4 T cells (left) and phenotypic marker expression on gated TRM1 and TRH cells (right). **b-c,** Flow cytometry plot representation of CD4^+^T-bet ZsGreen^+^ cells within TRM1 gate **(b)** and CD4^+^Bcl6 RFP^+^ cells within TRH gate **(c).** NP-specific TRM1 and TRH are shown on the left for comparison (n=4, representing 2 experiments).

**Supplementary Fig. 7.**
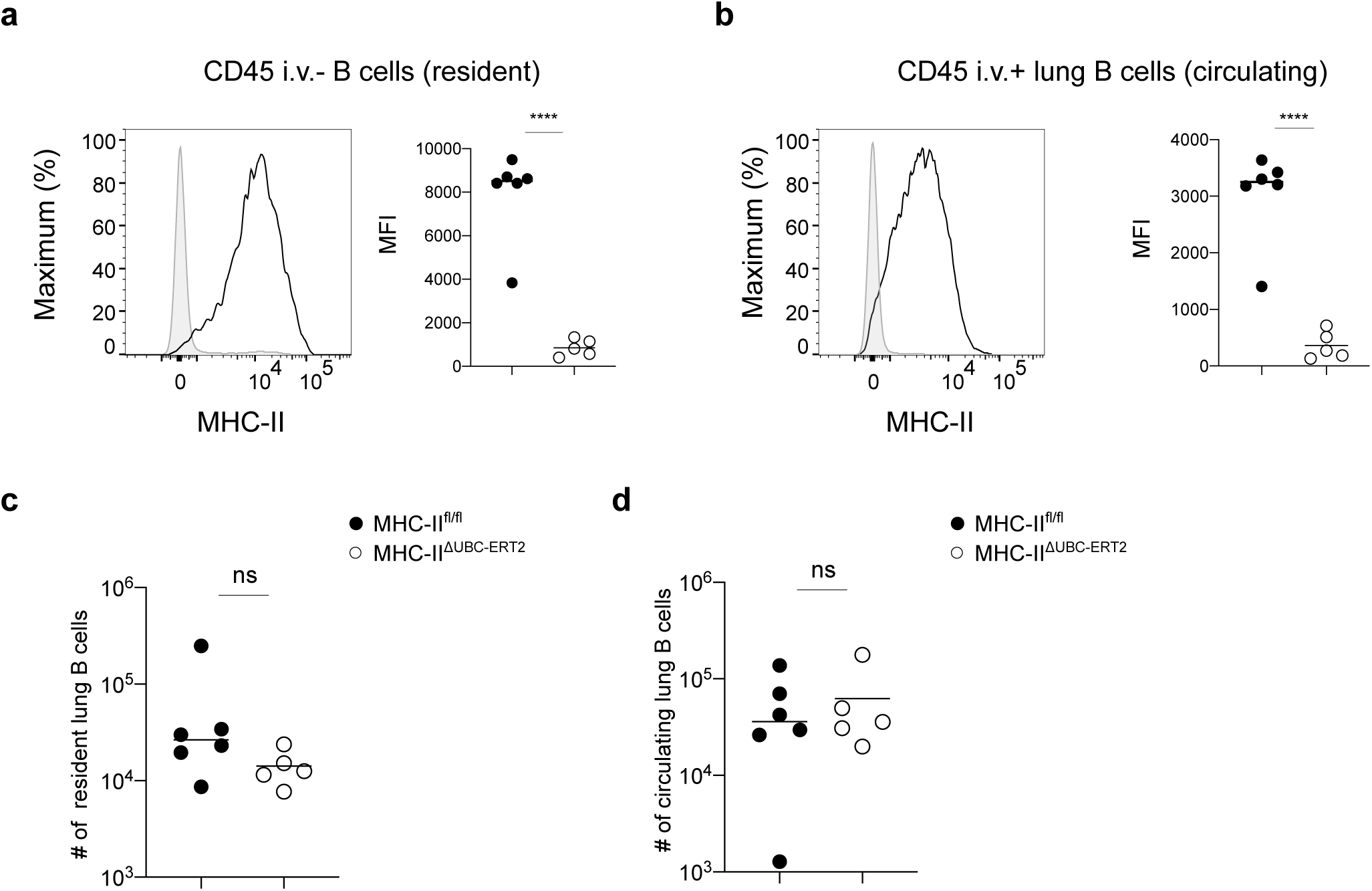
Maintenance of TRH cells requires antigen presentation. **a-b,** Histograms (left) and quantification (right) of MHC-II expression on iv^-^ **(a)** and iv^+^ B cells **(b)** upon inducible deletion of MHC-II. **c-d,** Total numbers of iv^-^ **(c)** and iv^+^ **(d)** lung B cells. Thin lines represent mean (n=5-6, representing 2 experiments). Significance was determined using unpaired Student’s t-test. P values are as follows: *P<0.05, **P<0.01, ***P<0.001, ****P<0.0001.

**Supplementary Fig. 8.**
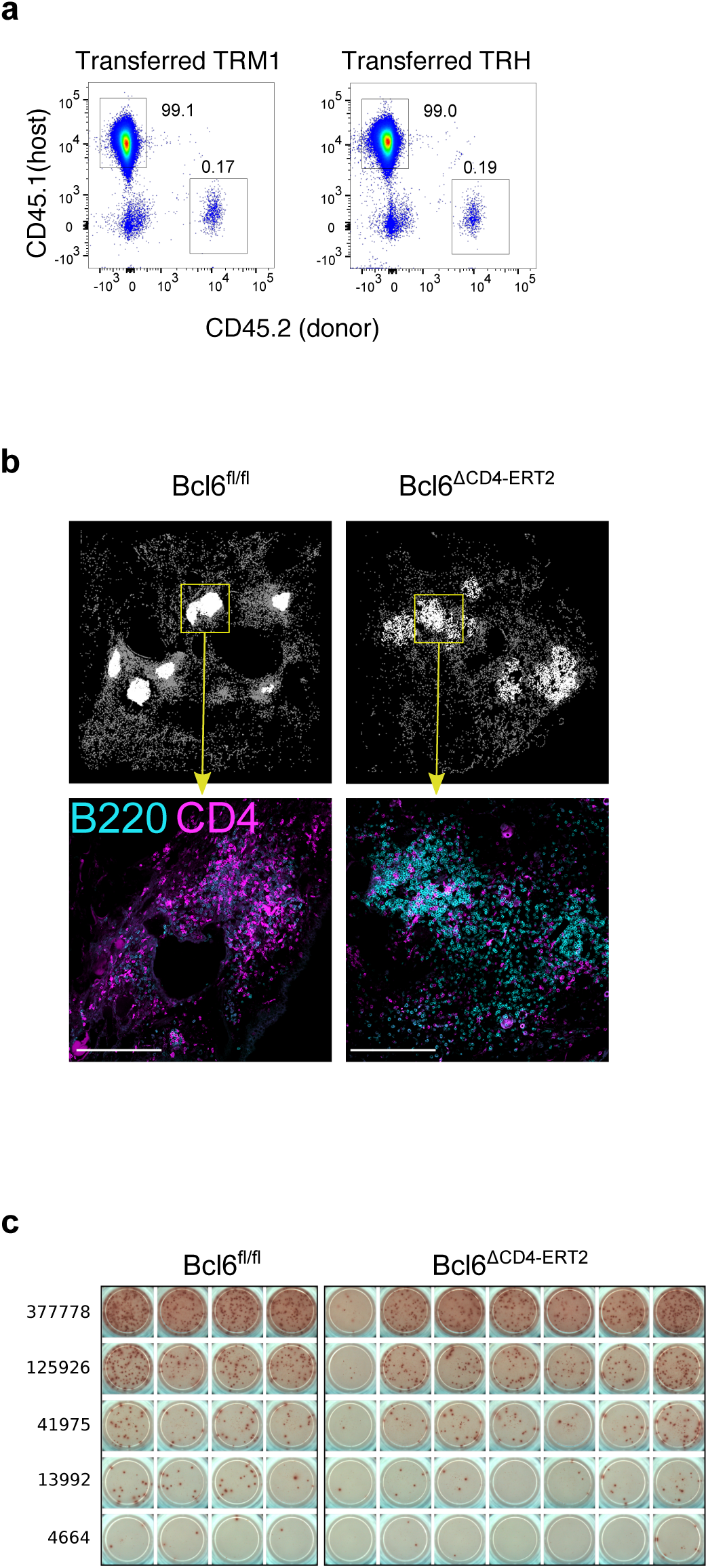
TRH cells contribute to protection during re-challenge. **a**, Flow cytometry plots representing proportion of resident CD4 T cells in donor vs host in TRM1 and TRH recipients. **b,** Spatial distribution of CD4 cell objects (top) quantified from immunofluorescence confocal images (bottom) from lungs of Bcl6^flox/flox^ and Bcl6Δ^CD4-ERT2^ mice, 30 days post-infection. Each dot represents a CD4+ T cell object, located in a B220+ area (white), or outside B220+ areas (grey). **c,** Representative pictures of ELISpot for NP-specific IgG ASC from lungs of Bcl6^fl/fl^ and Bcl6^ΔCD4-ERT2^ mice. Data in **(a)** represents 2 experiments, n=3 and **(b)** represents 2 experiments, n=4-7.

